# An Innovative Framework for Heart Sound Classification Integrating Adaptive Fuzzy Rank-Based Ensemble of Transfer Learning Models with Multi-Dimensional Features

**DOI:** 10.1101/2025.09.07.674773

**Authors:** Shuvashis Sarker, Faika Fairuj Preotee, Tashreef Muhammad, Shamim Akhter

## Abstract

Heart sound identification for diagnosing cardiovascular diseases is a complex challenge due to the intricate, spectro-temporal characteristics of conditions like Aortic Stenosis, Mitral Regurgitation, and multi-valvular disorders. Conventional methods frequently inadequately encompass the complete range of diagnostic characteristics, depending on fragmented or simplistic approaches that lack generalizability across varied patient demographics, clinical settings, and recording circumstances. To address these limitations, we offer an innovative, multi-dimensional feature fusion framework that integrates Wavelet Scattering Transform (WST) and Mel-Frequency Cepstral Coefficients (MFCC) to capture both temporal stability and perceptually optimal frequency patterns from cardiac sounds. This method is further refined by an adaptive fuzzy rank-based ensemble technique utilizing the Gompertz function, which dynamically modifies model weights according to confidence metrics, hence assuring more dependable and precise predictions amidst fluctuating clinical uncertainty. We meticulously assess our model utilizing eight advanced fine-tuned transfer learning architectures across four feature extraction techniques (WST, MFCC, STFT, Multi-Dimensional) on the clinically validated BUET Multi-disease Heart Sound (BMD-HS) dataset. This dataset comprises 864 phonocardiogram recordings from 108 participants with echocardiographically confirmed diagnoses. The multi-dimensional feature fusion method attains 97% accuracy with EfficientNetB2, whereas the adaptive fuzzy ensemble strategy achieves 98% accuracy, surpassing both individual models and conventional ensemble methods. Moreover, Explainable AI with Audio-LIME offers transparent, clinically interpretable insights, achieving fidelity scores surpassing 0.85 and clinical relevance ratings exceeding 90%, facilitating the identification of critical time-frequency regions that hold diagnostic significance. This system establishes a novel benchmark for heart sound classification, enhances the differentiation of intricate valvular disorders, and provides dependable, evidence-based decision support for cardiovascular diagnosis in clinical settings. Our research illustrates the capability for resilient, scalable, and interpretable heart sound classification systems that can improve clinical decision-making and promote the integration of automated diagnostic tools in healthcare.

## Introduction

The human heart, commonly known as the primary pump of the body, is an extraordinary muscular organ that is tasked with preserving blood circulation and sustaining life. It facilitates the delivery of oxygen and vital nutrients to tissues by regular contractions, simultaneously eliminating carbon dioxide and metabolic waste. The scientific comprehension of this organ commenced with the identification of blood circulation and the heart’s crucial function in propelling it [1, 2]. Subsequent research demonstrated that cardiac output is governed by both intrinsic heart performance and the requirements of bodily tissues, underscoring the intricacy of cardiovascular regulation [3]. Research on the contractile mechanics of the myocardium has enhanced our understanding of the heart’s response to physiological stress and pharmacological therapies [4]. In recent decades, extensive epidemiological studies [5, 6] have demonstrated that heart disease continues to be a primary cause of mortality worldwide, highlighting the importance of lifestyle choices and preventive measures. These fundamental discoveries persist in shaping both scientific and clinical methodologies about cardiac health.

Heart sounds arise from the mechanical activity of the heart, mostly due to the closing of cardiac valves and the flow of blood. Table 1 shows the characteristics of heart sound from S1 to S4 [7]. Murmurs, extended sounds resulting from turbulent flow due to valve stenosis, regurgitation, or septal anomalies, offer critical diagnostic information. Phonocardiography, the analysis of cardiac sounds, has progressed markedly with the use of digital recording technology and computational methods. Phonocardiogram (PCG) [8] signals, which capture acoustic emissions of cardiac activity using sensitive microphones, provide a high-fidelity representation of heart sounds and murmurs, facilitating advanced signal analysis for diagnostic purposes. Electrocardiogram (ECG) data offer electrical insights into the cardiac cycle and are frequently utilized alongside PCG recordings to investigate temporal relationships between electrical and mechanical events in the heart. The relationship between acoustic and electrical signals is essential in modern cardiology, especially in machine learning applications.

**Table 1.** Characteristics of Heart Sounds (S1–S4) [9].

| Heart Sound | Cause | Timing in Cardiac Cycle | Clinical Significance |
| --- | --- | --- | --- |
| S1 | Closure of mitral and tricuspid valves | Beginning of systole | Normal sound |
| S2 | Closure of aortic and pulmonary valves | End of systole | Normal sound |
| S3 | Extra sound after S2 | Early diastole | Ventricular Dysfunction |
| S4 | Extra sound before S1 | Late diastole | Stiff Ventricles |

Despite progress in digital phonocardiography and machine learning for heart sound analysis, significant research deficiencies remain, hindering the clinical efficacy of automated cardiac diagnostic systems. Contemporary methodologies depend on isolated feature extraction methods [10,11] such as MFCC or rudimentary time-frequency analysis, which inadequately capture the intricate spectro-temporal characteristics essential for differentiating delicate cardiovascular illnesses. Moreover, these approaches fail to accommodate the variability in heart sounds across different patient demographics, recording conditions, and disease states, hence diminishing their generalizability from research to clinical applications. Ensemble methods [12] in heart sound classification frequently rely on simplistic voting mechanisms or static weight distributions, failing to leverage the complementary advantages of deep learning architectures. Additionally, the absence of multi-dimensional feature fusion algorithms [13] and adaptive ensemble techniques undermine predictive accuracy, particularly across diverse cardiovascular conditions, thereby obstructing the advancement of clinically applicable tools that correspond to cardiologists’ decision-making processes.

The key innovations of our approach include:

i. **Systematic Integration of Multi-Dimensional Features:** Combines Wavelet Scattering Transform (WST) and Mel-Frequency Cepstral Coefficients (MFCC) for a robust, multi-dimensional feature set in heart sound classification.
ii. **Fine-Tuned Transfer Learning Architectures:** Evaluates eight transfer learning architectures (VGG, ResNet, DenseNet, EfficientNet, Inception, Xception, MobileNet) to identify the best-performing models.
iii. **Adaptive Fuzzy Rank-Based Ensemble Method:** Introduces an adaptive fuzzy rank-based ensemble method using the Gompertz function, dynamically adjusting model weights to achieve optimal results based on confidence metrics.
iv. **Multi-Dimensional Feature Visualization and Interpretability:** Develops a framework using Audio-LIME to visualize and interpret multi-dimensional features for clinical insights.

The unique approaches presented in this research address existing deficiencies in heart sound analysis by providing a more robust, flexible, and clinically pertinent approach. We improve diagnostic accuracy by integrating multi-dimensional feature fusion with an adaptive ensemble model, therefore addressing the intrinsic unpredictability in heart sound data.

### Related Works

The realm of automated heart sound classification has experienced significant evolution and advancements due to the systematic implementation of advanced machine learning and deep learning techniques, with extensive research persistently investigating innovative methods to improve diagnostic accuracy, clinical reliability, and practical applicability in cardiovascular disease detection. The analysis of heart sounds, as thoroughly detailed in Table 2, involves a wide range of methodological approaches. Researchers are exploring various advanced combinations of preprocessing techniques, feature extraction methods, and classification algorithms to effectively tackle the complex challenges associated with PCG analysis for identifying cardiovascular pathologies. These thorough investigations encompass a broad array of datasets, including the widely-utilized PhysioNet/CinC 2016 challenge dataset [10, 11, 14, 15], the ICBHI 2017 respiratory sound database [16, 17], Urban Sound collections [18, 19], and various proprietary clinical datasets [20, 21], each exhibiting distinct characteristics and challenges that necessitate customized analytical methodologies. The preprocessing strategies utilized in these studies exhibit significant diversity, encompassing basic signal processing techniques such as signal segmentation [10, 22], resampling, and noise filtering with Butterworth bandpass filters [11, 23],, as well as more advanced methodologies including intricate data augmentation techniques [21,24], pitch shifting, time stretching [16,21], normalization procedures [18,20,25], and specialized cardiac cycle segmentation methods [14, 26]. Feature extraction methodologies have significantly evolved, moving from traditional signal processing techniques like Mel-Frequency Cepstral Coefficients (MFCC) [11, 16, 17, 20, 21], which yield perceptually relevant frequency domain representations, and Wavelet Scattering Transform (WST) [10, 27–29], which captures multi-scale temporal invariant features, to more advanced spectro-temporal representations such as Short-Time Fourier Transform (STFT) [18, 21, 25], Constant-Q Transform (CQT) [17], mel spectrograms [16, 30, 31], chromagrams [16, 18], and hybrid feature fusion approaches [23] that aim to utilize complementary information from various feature domains. Classification methodologies have undergone a paradigmatic transformation from traditional machine learning algorithms, such as Support Vector Machines (SVM) [11, 18, 19], Random Forests [15], K-Nearest Neighbors (KNN) and Decision Trees [11, 15, 24], which depend on manually crafted features and established statistical learning principles, to more intricate and potent deep learning architectures. These include Convolutional Neural Networks (CNNs) for spatial pattern recognition [10, 16, 25, 30], Long Short-Term Memory (LSTM) networks and Bidirectional LSTM (BiLSTM) for temporal sequence modeling [18, 25, 31], Gated Recurrent Units (GRUs) for efficient sequential processing [20, 31], ResNet architectures for deep residual learning [14,29,32], transformer-based models for attention-driven feature learning [22,28], and advanced ensemble methods that integrate multiple models to enhance predictive performance [10, 21, 30].

**Table 2.**
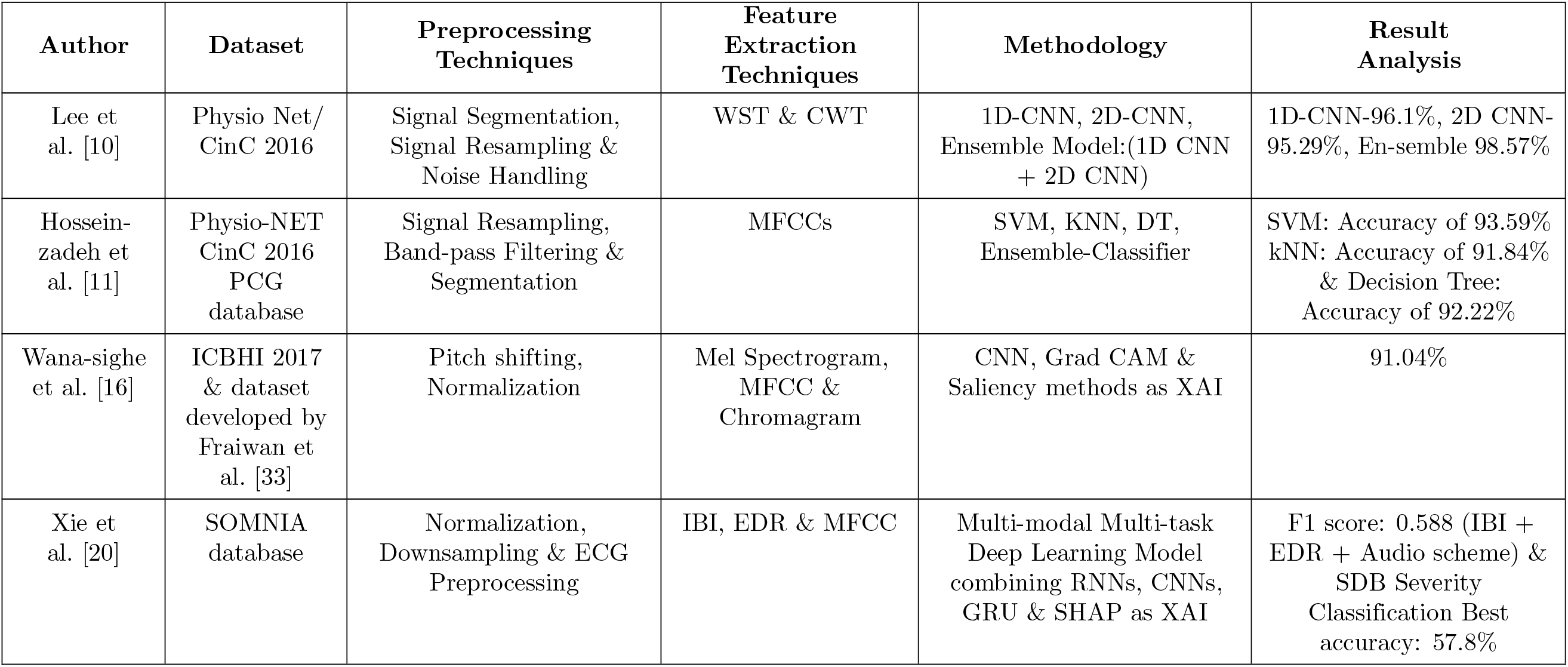
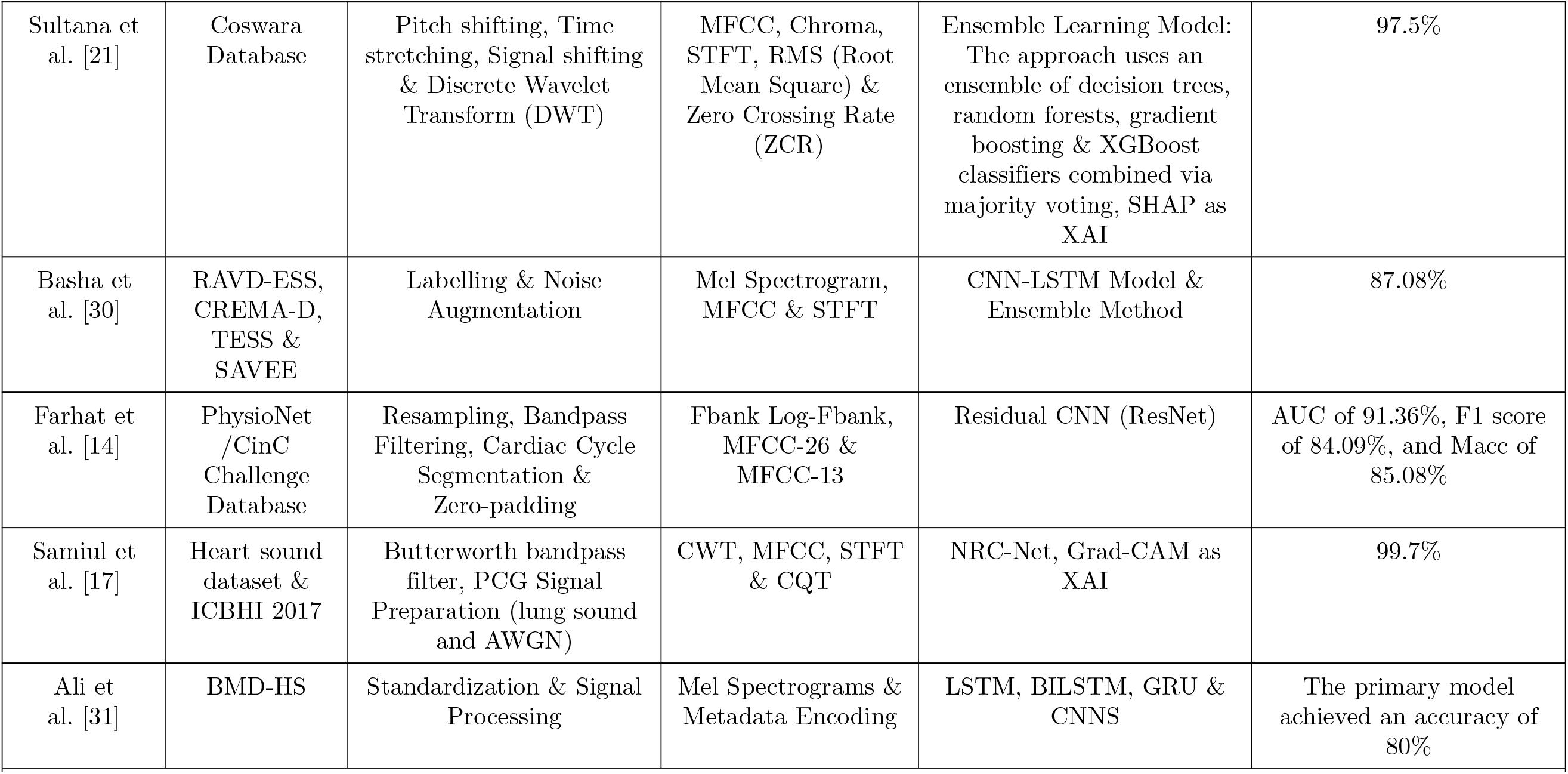
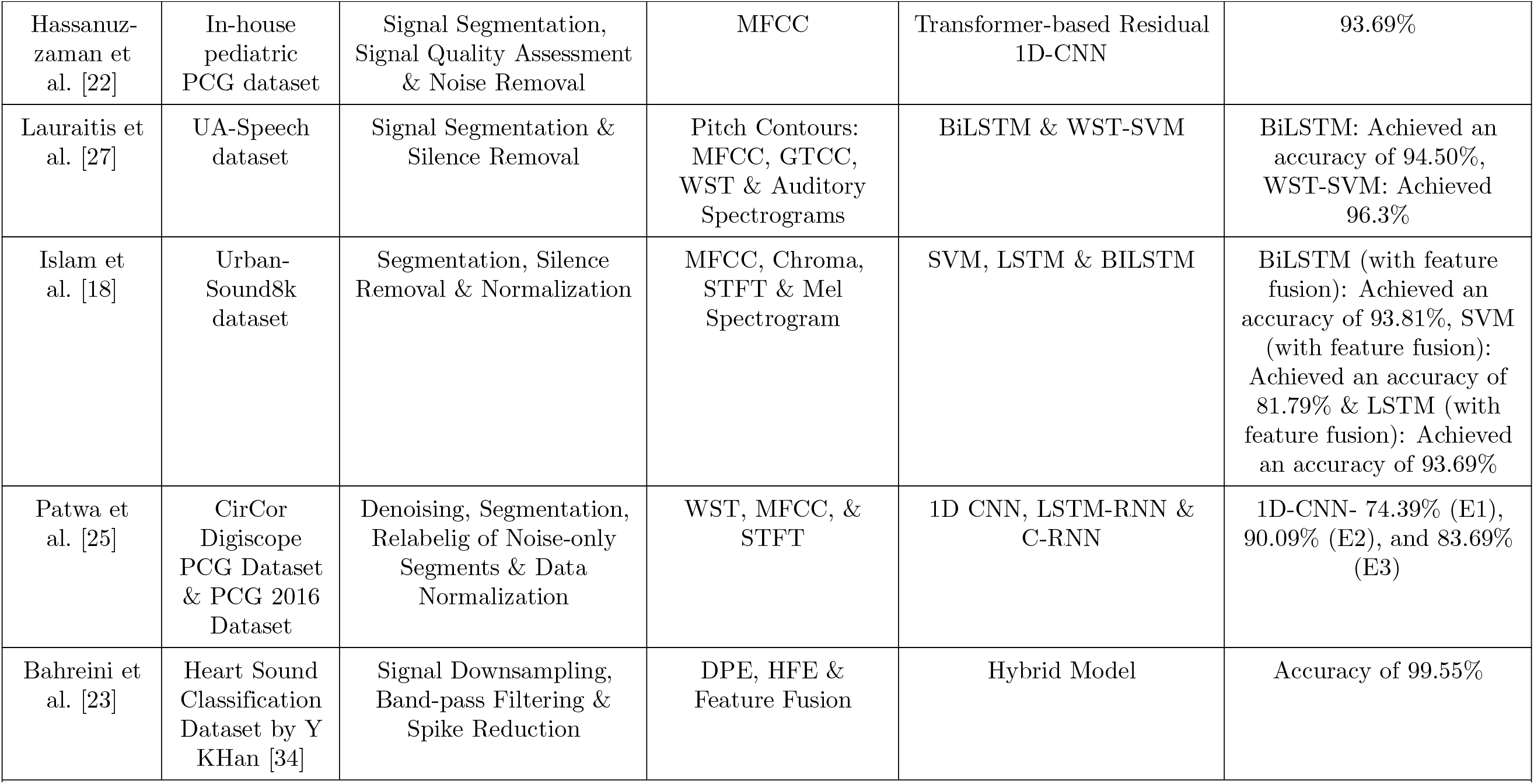
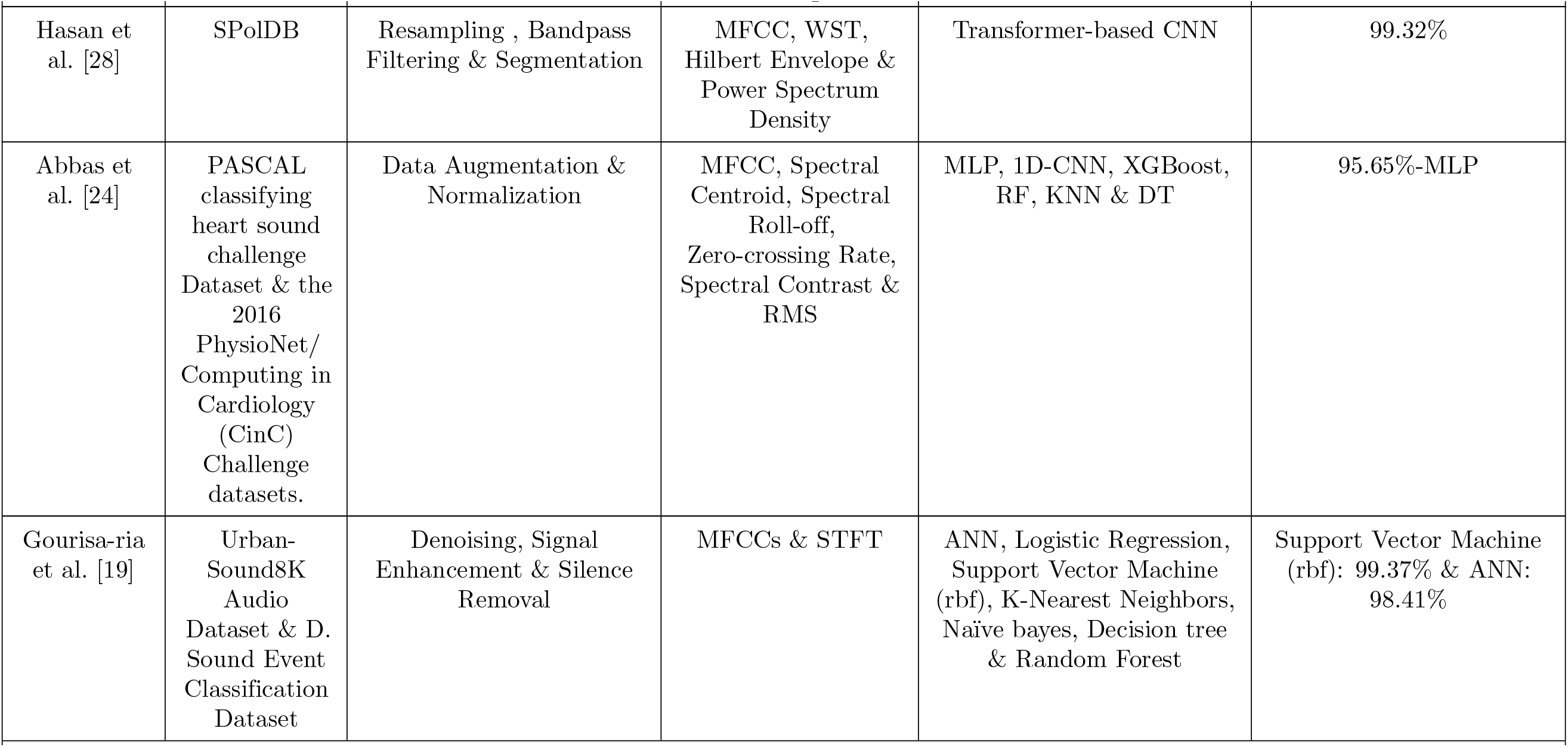
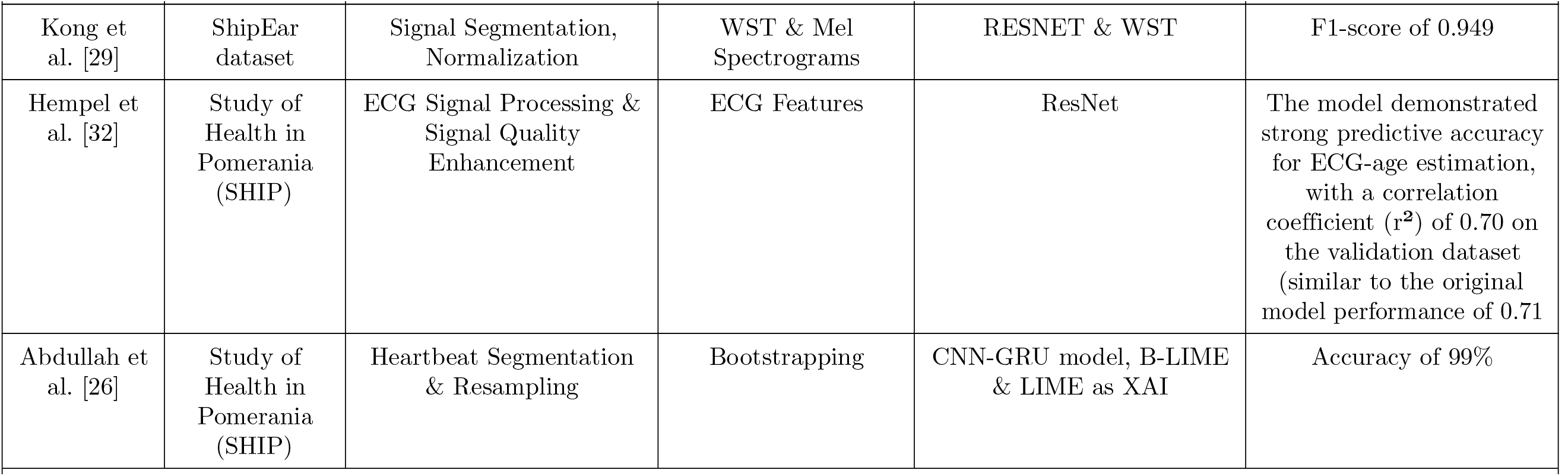
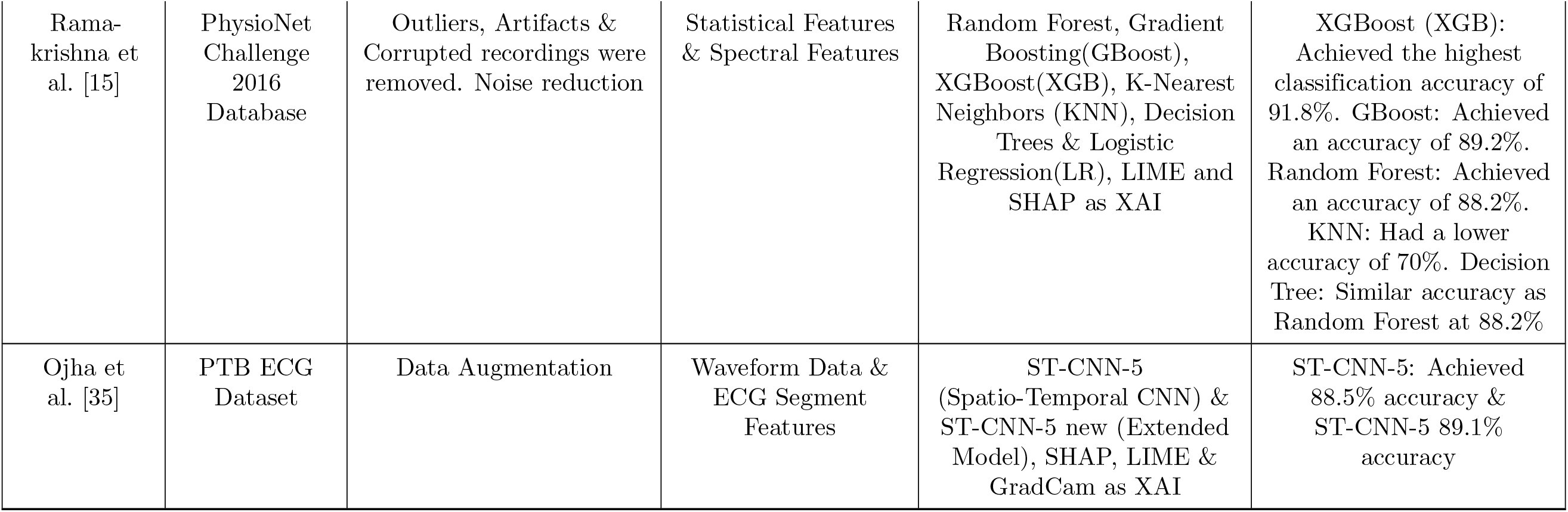
Comparative Analysis of Previous Studies.

This research presents a comprehensive framework to address key limitations in heart sound classification systems, enhancing clinical viability and robustness. The multi-dimensional feature integration challenge is addressed by combining Wavelet Scattering Transform and Mel-Frequency Cepstral Coefficients, resulting in a cohesive spectro-temporal feature space that captures complementary temporal stability and perceptually optimized frequency representations. To bridge the explainability gap, Audio-LIME is employed for multi-dimensional feature interpretation, offering clinically relevant insights that enable healthcare professionals to understand the influence of various attributes on diagnostic decisions, thereby fostering clinical trust and ensuring regulatory compliance. The limitations of static ensemble methods are overcome by an adaptive fuzzy rank-based ensemble approach, which uses the Gompertz function for dynamic confidence-based weighting. This system adjusts intelligently to fluctuations in uncertainty and clinical data, ensuring consistent performance in diverse real-world settings. Architectural limitations are mitigated by the fine-tuning of eight advanced transfer learning architectures, identifying the most effective models for various clinical conditions and recording environments. Finally, clinical deployment reliability is assured through stringent validation protocols and benchmarking, ensuring high diagnostic accuracy and meeting the rigorous standards necessary for healthcare implementation, thus enabling practical adoption and regulatory approval in real-world medical settings.

### Dataset

The BUET Multi-disease Heart Sound (BMD-HS) dataset [31] is a clinically validated repository of PCGs that is publicly accessible [36]. It includes 864 recordings from 108 individuals (87 patients with valvular heart disorders and 21 healthy controls). The dataset encompasses four common valvular heart diseases: Aortic Stenosis (AS), Aortic Regurgitation (AR), Mitral Stenosis (MS), and Mitral Regurgitation (MR). These conditions substantially contribute to global cardiovascular mortality, with aortic valve disorders responsible for 61% and mitral valve diseases for 15% of deaths related to valvular heart disease [37]. Figure 1, Figure 2, and Figure 4 depict representative PCG waveforms that exhibit unique heart sound patterns corresponding to each situation. The data collection was conducted at the National Institute of Cardiovascular Diseases (NICVD) in Dhaka, Bangladesh, utilizing the 3M™Littmann^®^ Model 3200 digital stethoscope with a 4 kHz sampling rate. Participants were recorded in both sitting and supine postures at four auscultation sites (mitral, tricuspid, pulmonic, aortic), resulting in eight 20-second recordings per individual. Recordings were transformed from.zsa to.wav format using 3M™Littmann^®^ StethAssist™, using quality control methods to ensure data integrity.

**Fig 1.**
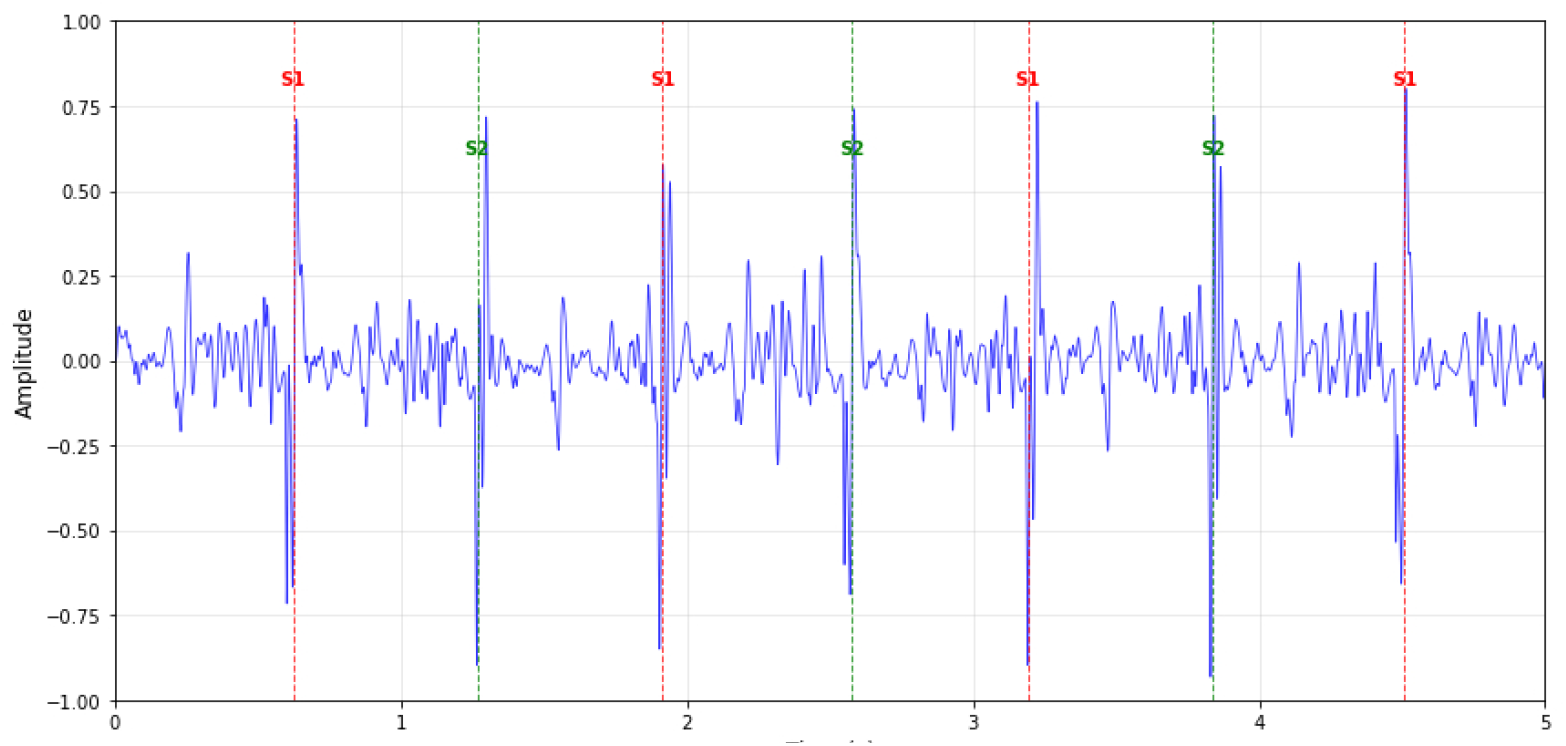
PCG Signal Segment of Diseased Heart (MR & AR) from BMD-HS Dataset with Bell Mode Filtering.

**Fig 2.**
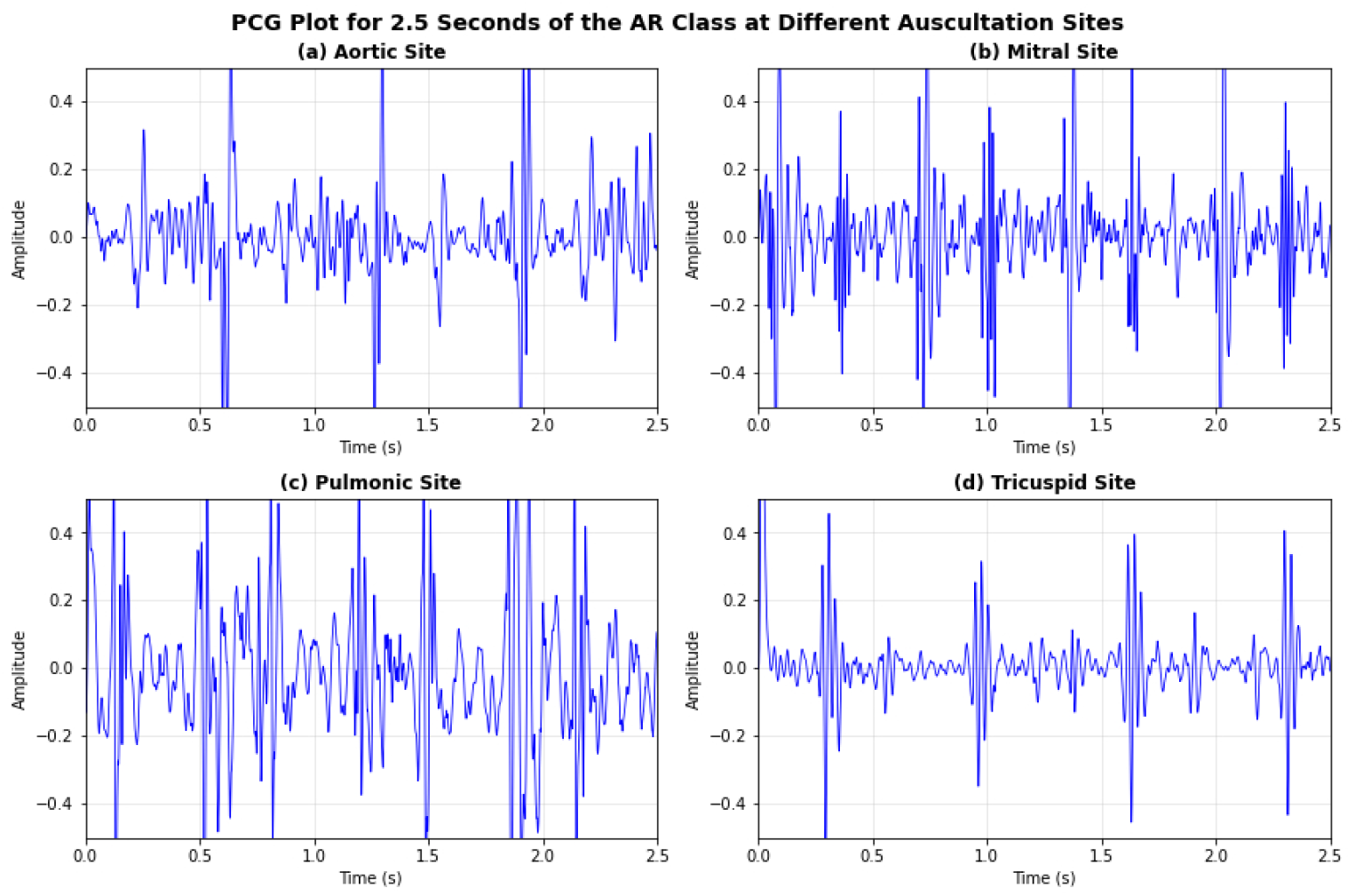
Multi-site PCG recordings showing waveforms from different auscultation sites (aortic, mitral, pulmonic, tricuspid).

Clinical validation entailed echocardiogram reports that corroborated all diagnoses, offering gold-standard verification and reliable ground truth labeling. The dataset consists of six categories: Normal (N=21), AS (N=8), AR (N=7), MR (N=11), MS (N=11), and Multi-disease (N=50), as illustrated in Figure 3. The significant multi-disease category fills a crucial void by encompassing concurrent multiple valve disorders, which are infrequently depicted in current datasets. An advanced multi-label annotation system records individual and numerous disease states alongside extensive demographic information (age, gender, smoking status, geographic area, disease severity). Environmental consistency was preserved by utilizing pre-recorded NICVD hospital background noise in laboratory settings, especially during COVID-19, hence providing uniform acoustic features throughout all recordings. The dataset illustrates the current epidemiology of valvular heart disease in Bangladesh and acts as a standard for automated heart sound classification and auscultation-based diagnostic instruments in resource-constrained healthcare environments.

**Fig 3.**
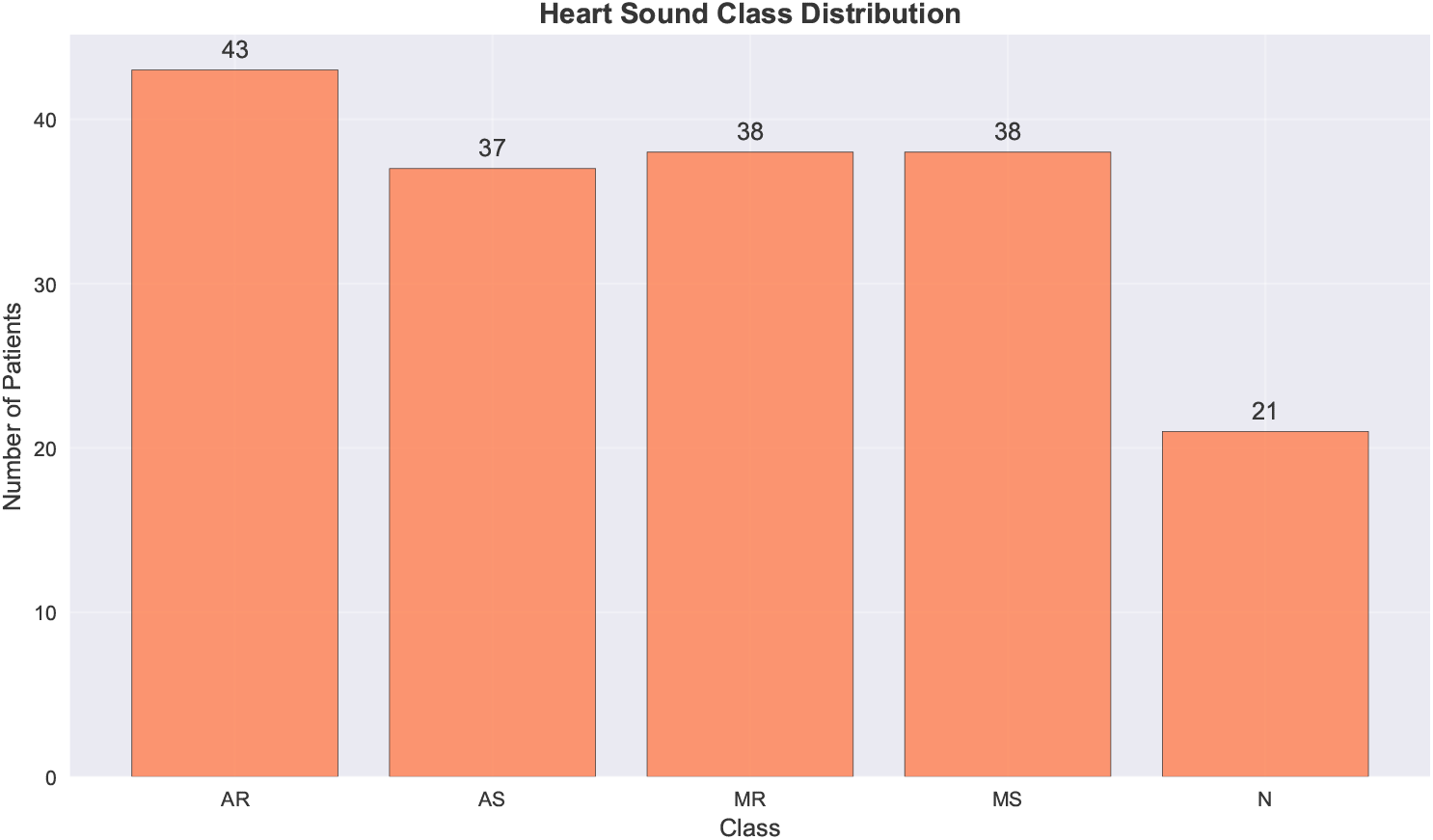
Class distribution showing the number of patients in each disease category.

**Fig 4.**
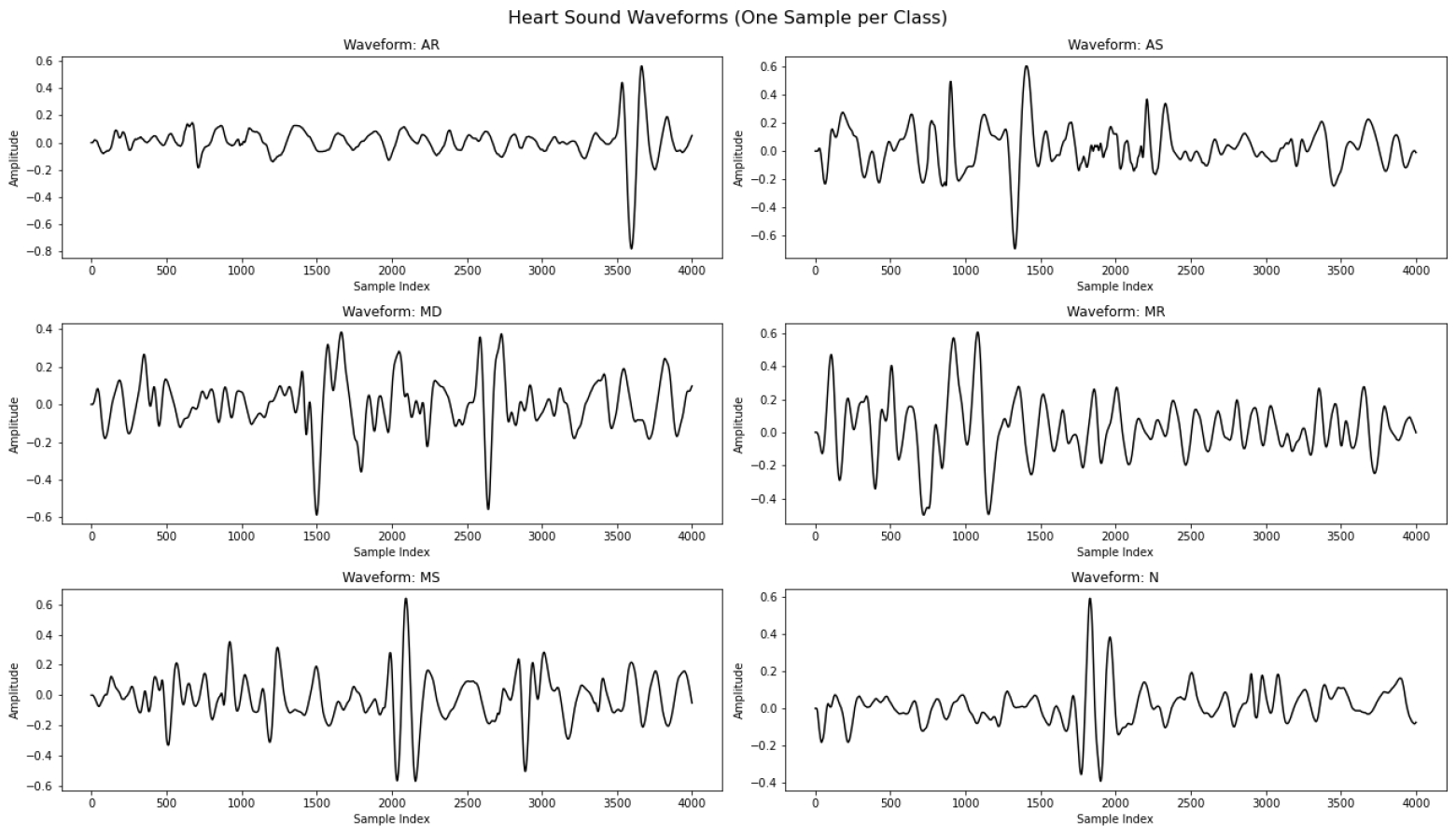
Representative waveforms showing one sample from each disease class (AR, AS, MR, MS, MD, N) to illustrate characteristic patterns.

### Proposed Methodology

Our proposed framework combines multi-dimensional feature fusion with an adaptive fuzzy rank-based ensemble approach for automated heart sound categorization, as depicted in Figure 5. The method analyzes PCG signals from the BMD-HS dataset via thorough data preprocessing, subsequently extracting WST, MFCC, STFT, and Multi-Dimensional features that encapsulate complimentary spectro-temporal attributes of heart diseases. Eight meticulously optimized transfer learning architectures are trained on these feature representations, and their predictions are amalgamated using our innovative Gompertz function-based fuzzy ensemble method, with explainable AI via AudioLIME offering clinically interpretable insights into the decision-making process.

**Fig 5.**
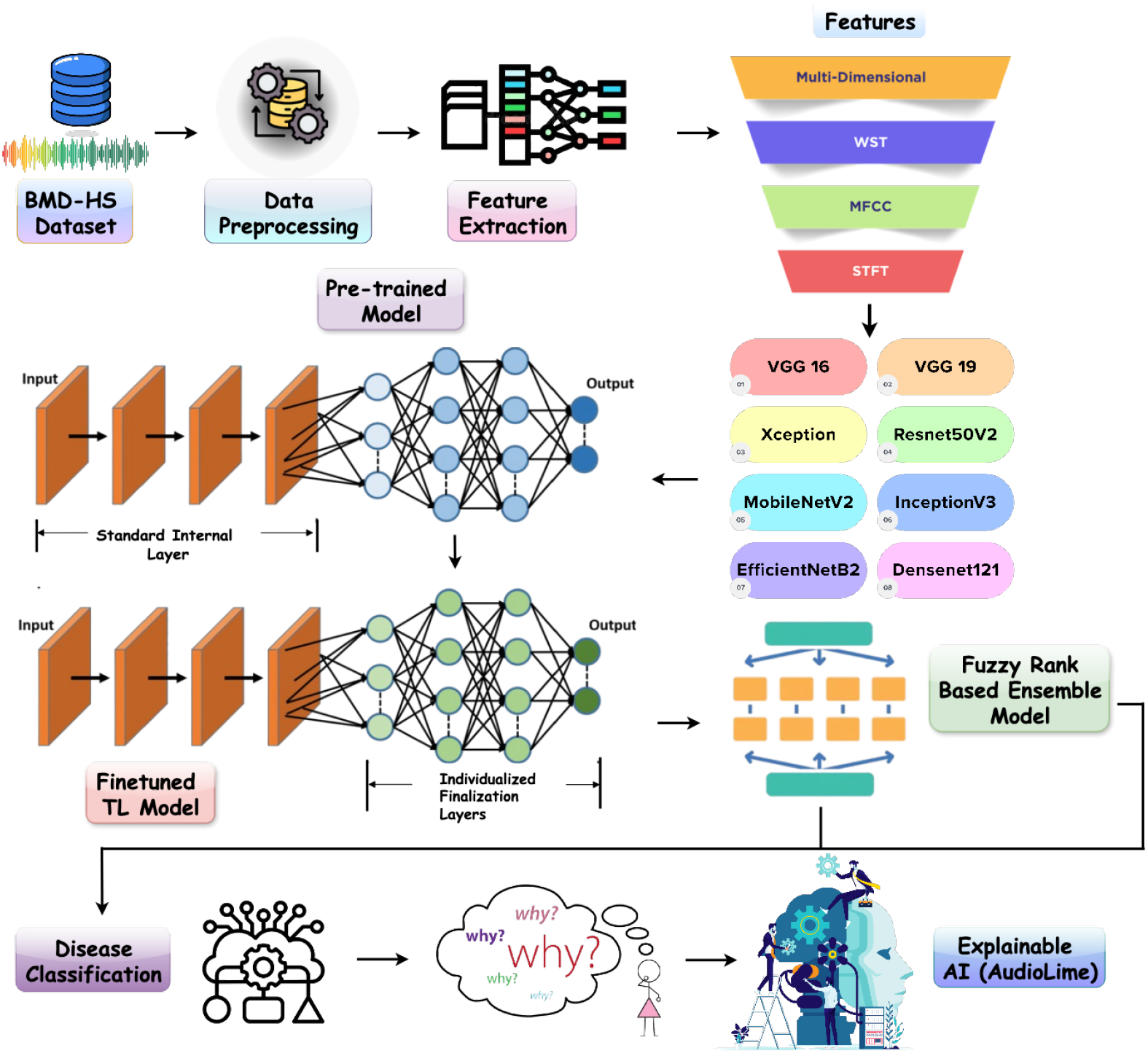
Proposed Methodology

### Data Preprocessing

Audio data preparation is essential for improving the efficacy of machine learning models in PCG analysis for cardiovascular disease identification. This section delineates a preparation workflow specifically designed for the BMD-HS dataset, tackling its distinct issues. Figure 6 illustrates the pipeline developed to enhance signal quality for effective feature extraction in later stages.

**Fig 6.**
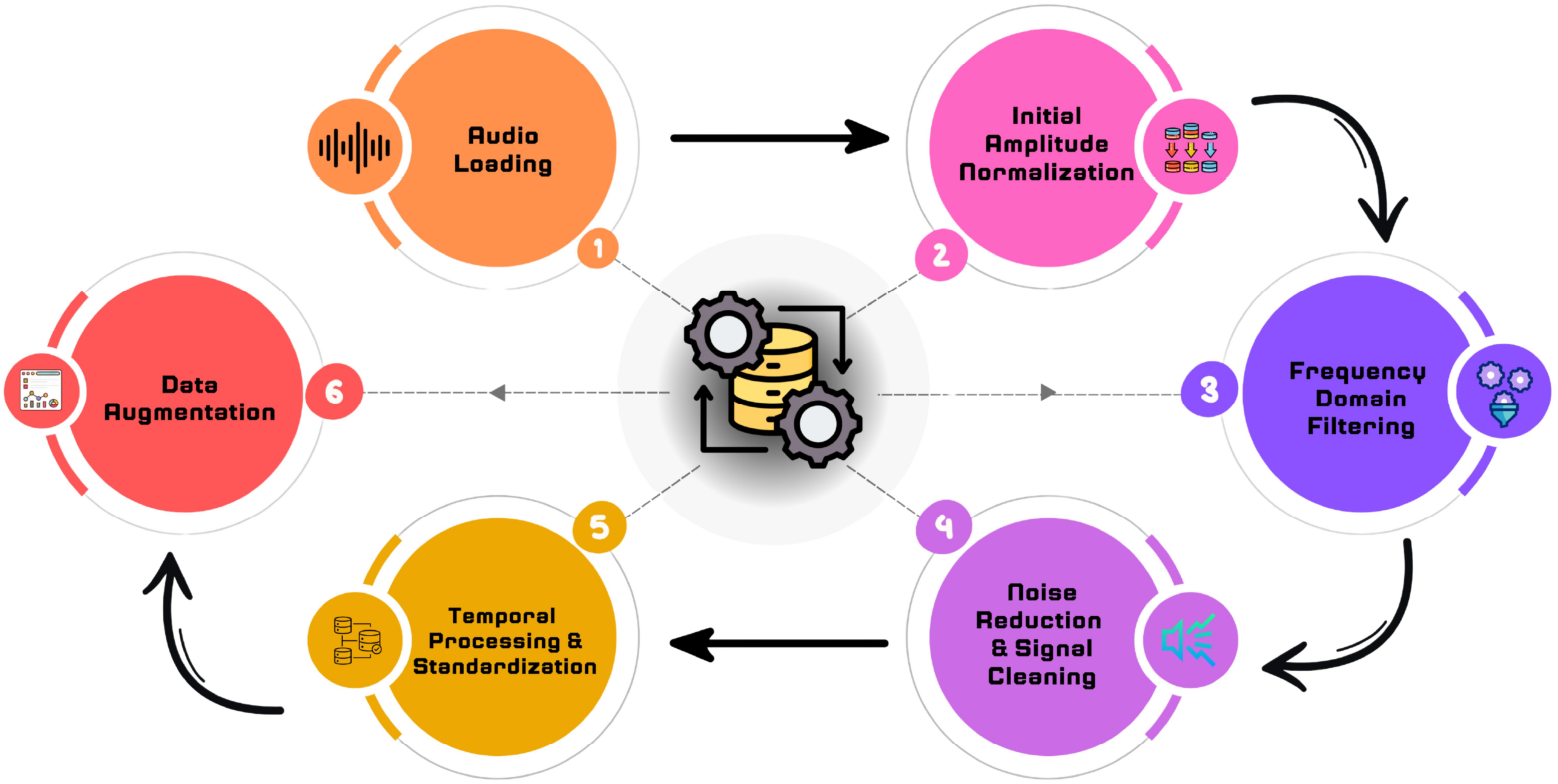
The workflow of data preprocessing steps for audio signals.

#### Audio Normalization

The BMD-HS dataset exhibits amplitude fluctuations in raw PCG recordings due to patient positioning, auscultation site variations, and individual physiological differences. Audio normalization is essential for PCG analysis to standardize signals from diverse clinical environments and patient populations, where inconsistent recording conditions significantly impact diagnostic accuracy. The preprocessing pipeline begins with audio loading, converting multi-channel recordings to mono format, followed by resampling to a uniform 4 kHz rate. This sampling rate effectively captures essential heart sound frequencies and harmonics up to 2 kHz according to the Nyquist theorem, while reducing computational complexity. The 4 kHz rate is optimal for valvular heart disorder analysis, as diagnostically significant spectral content for aortic stenosis, aortic regurgitation, mitral stenosis, and mitral regurgitation occurs below 1 kHz. Higher sampling rates introduce unnecessary processing overhead and potential high-frequency artifacts that compromise classification performance [38–41]. Amplitude normalization standardizes signal range across recordings using:

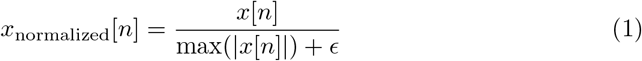

where *x*[*n*] represents the discrete-time audio signal, max(|*x*[*n*|]) is the maximum absolute amplitude, and *ϵ* = −1 × 10^−9^ prevents division by zero for silent segments [42]. This normalization scales all signals to the [−1, 1] range, ensuring consistent amplitude levels across the 108 subjects and preventing amplitude variations from dominating the feature space. By implementing this audio normalization pipeline, PCG analysis achieves improved signal-to-noise ratio, enhanced amplitude consistency, and reduced computational overhead, thereby significantly improving classification accuracy and generalizability of automated heart sound analysis systems.

#### Frequency Domain Filtering

The BMD-HS dataset contains frequency components outside the diagnostically relevant range that may compromise cardiovascular disease classification. Clinical PCG recordings include low-frequency ambient noise, patient movement artifacts, and high-frequency interference from digital stethoscopes and wireless systems. These undesirable spectral components increase noise, computational costs, and may cause erroneous feature learning in deep neural networks.

A fourth-order Butterworth bandpass filter preserves frequencies within the 20-900 Hz range, based on cardiovascular research and valve disease classification criteria. The Butterworth filter provides superior flat passband response and smooth transition properties, minimizing distortion of essential frequencies while efficiently attenuating unwanted components. The filter transfer function is:

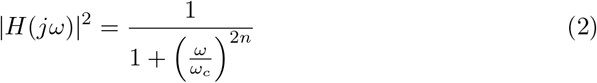

where *ω* is the angular frequency, *ω*_*c*_ is the cutoff frequency, and *n* = 4 is the filter order [43]. The normalized frequency parameters are:

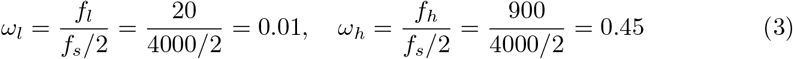

where *f*_*l*_ = 20 Hz and *f*_*h*_ = 900 Hz represent the cutoff frequencies, and *f*_*s*_ = 4000 Hz is the sampling frequency [44].

Figure 7 illustrates the efficacy of bandpass filtering by evaluating raw and filtered heart sound waveforms across various illness classifications.

**Fig 7.**
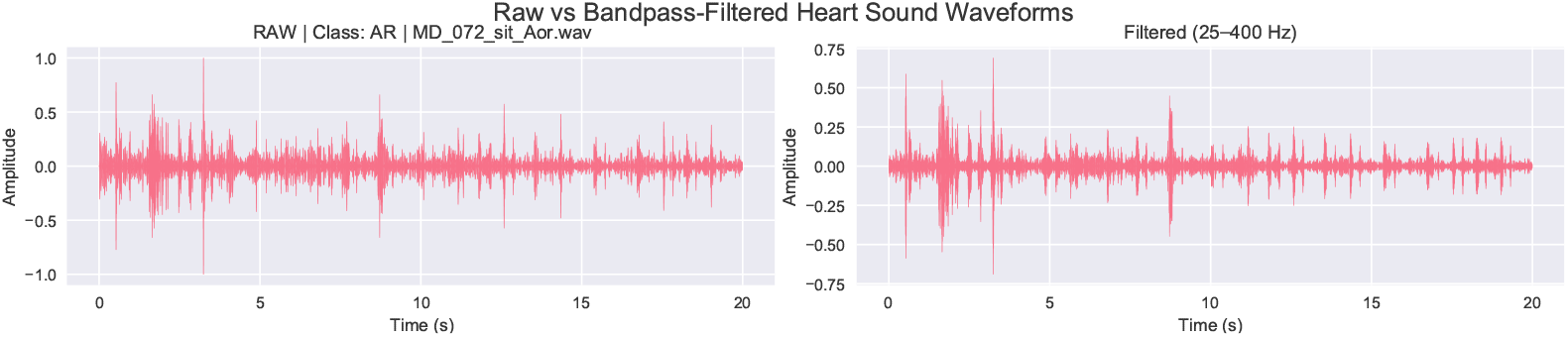
Comparison of raw and bandpass-filtered heart sound waveforms (25–400 Hz) across different disease classes.

The 20-900 Hz range encompasses all diagnostically relevant cardiac acoustic events while eliminating interference. Components below 20 Hz include environmental rumbling and mechanical vibrations, while components above 900 Hz comprise aliasing artifacts and electronic noise that disrupt classification systems. The filtering effectively eliminates low-frequency drift and high-frequency noise, improving S1 and S2 sound clarity and making murmurs more discernible. By implementing this frequency filtering approach, PCG analysis achieves enhanced signal clarity, improved feature extraction reliability, and reduced computational complexity, significantly improving diagnostic accuracy and robustness of automated heart sound classification systems.

#### Noise Reduction and Signal Cleaning

Clinical PCG recordings from the BMD-HS dataset contain various noise sources including hospital ambient noise, patient movement, respiratory sounds, and digital processing artifacts that compromise cardiovascular disease classification accuracy. The multisite recording approach (aortic, mitral, tricuspid, pulmonic) introduces diverse noise characteristics requiring systematic reduction.

An adaptive noise gate reduces low-amplitude background noise using a dynamically calculated threshold:

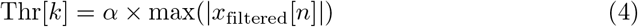

where *α* = 0.005 is the adaptive threshold coefficient, *x*_filtered_[*n*] is the bandpass-filtered signal, and *k* indexes individual recordings [45]. Audio samples below this threshold are attenuated:

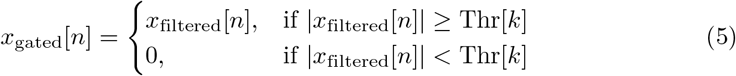

Numerical artifacts from floating-point computations and digital hardware are corrected using:

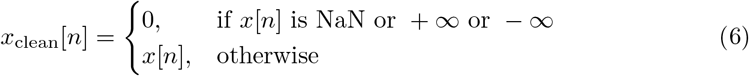

Impulse noise suppression employs median filtering with kernel half-width *k* = 2:

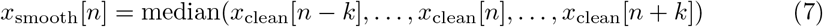

This preserves signal edges and transient features while removing outliers and spikes, maintaining S1 and S2 characteristics essential for valvular disease classification [46].

Final renormalization ensures consistent amplitude levels:

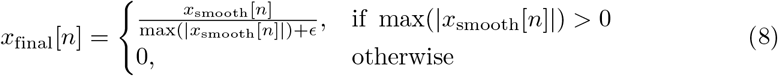

where *ϵ* = 1 × 10^−9^ provides numerical stability. Figure 8 demonstrates effective noise reduction through symmetric amplitude distributions, confirming well-normalized signals without clipping and validating preprocessing efficacy. This comprehensive noise reduction pipeline significantly improves signal quality, feature extraction accuracy, and reduces computational artifacts, enhancing the reliability of automated cardiovascular disease detection systems.

**Fig 8.**
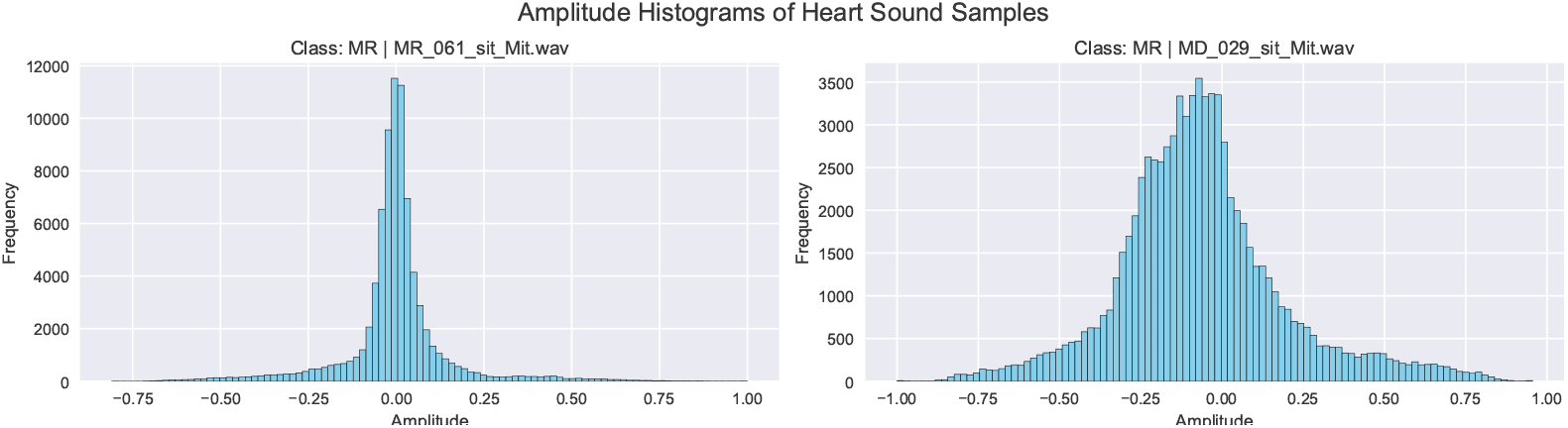
Amplitude histograms of heart sound samples illustrating the effectiveness of noise reduction and signal cleaning.

**Fig 9.**
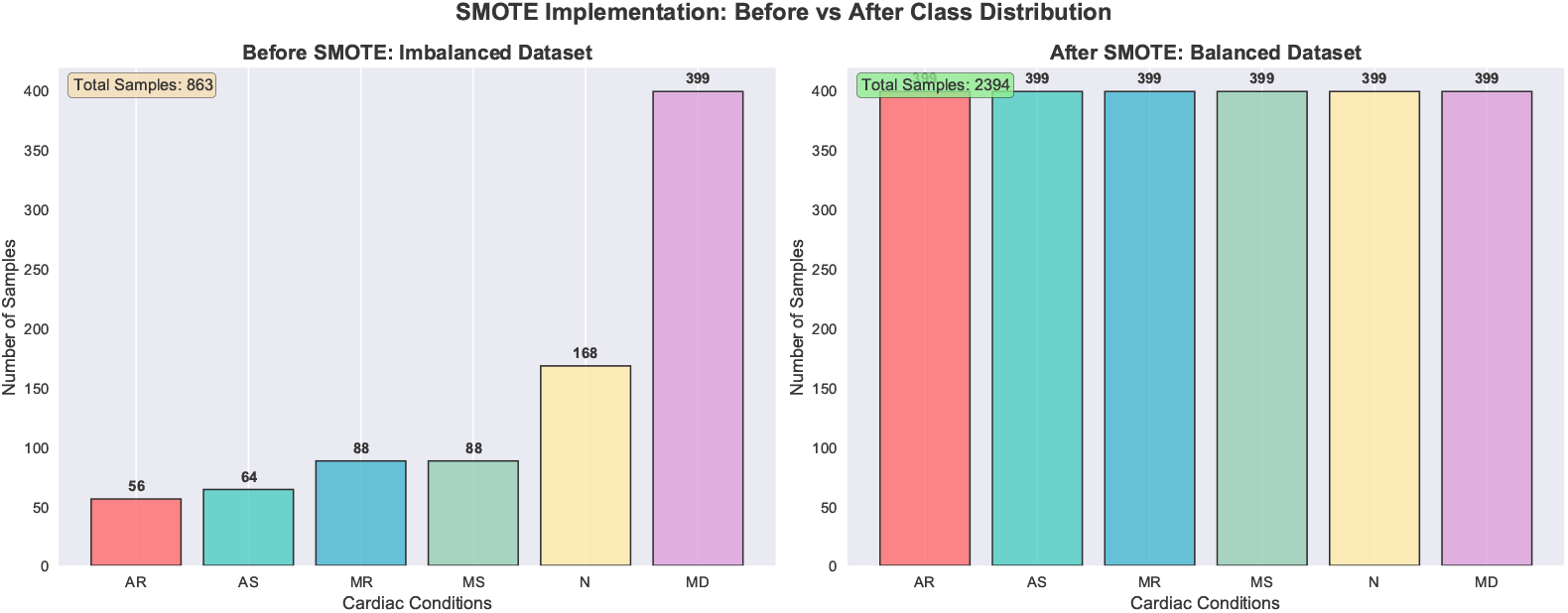
Class distribution before and after SMOTE application for balancing the dataset.

#### Temporal Processing and Standardization

The BMD-HS dataset exhibits temporal variability due to heart rate differences, recording timing variations, and silence intervals, complicating batch processing and model training. Temporal standardization ensures uniform signal duration and eliminates non-informative segments that introduce bias in automated cardiac assessment systems.

Trailing silence is removed using a detection technique that identifies the last sample containing cardiac data:

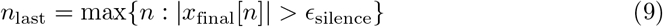

where *ϵ*_silence_ = 1 × 10^−6^ represents the silence threshold [47]. The signal is trimmed to the last informative sample:

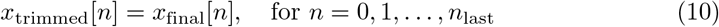

Adaptive padding and truncation normalize all recordings to 80,000 samples (20 seconds at 4 kHz), spanning 15-25 cardiac cycles for reliable valvular disease classification. For shorter recordings, zero-padding preserves temporal structure:

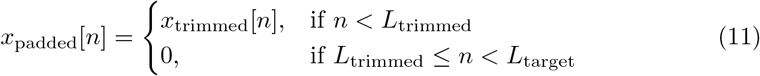

For longer recordings, truncation preserves initial segments containing highest quality cardiac information:

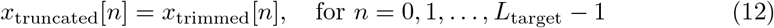

This standardization ensures consistent 80,000-sample inputs while maintaining cardiac timing relationships, providing uniform data structure for feature extraction techniques and enhancing machine learning model stability and performance [48].

#### Synthetic Minority Over-sampling Technique (SMOTE)

Data augmentation was utilized to expand the dataset by generating altered replicas of the original data, hence improving the generalization of machine learning models, especially for imbalanced datasets. The Synthetic Minority Over-sampling Technique (SMOTE) [49] was utilized to rectify class imbalance in the BMD-HS dataset. SMOTE creates synthetic samples by interpolation of existing data, thereby assuring that the generated data appropriately reflects minority classes. The SMOTE algorithm was executed by initially determining the *k*-nearest neighbors for each sample of the minority class by the use of Euclidean distance.

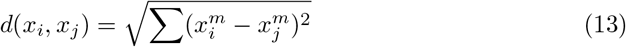

where 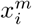 and 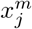 represent the m-th features of samples *x*_*i*_ and *x*_*j*_. Synthetic samples were created by generating new data points along line segments connecting original samples *x*_*i*_ and neighbors *x*_*j*_ using:

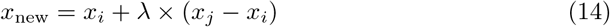

where *λ* is a random number between 0 and 1, ensuring synthetic samples lie within the range of original data and neighbors [50]. SMOTE was utilized on the BMD-HS dataset, which comprises 863 samples distributed among six categories: Aortic Regurgitation (AR), Aortic Stenosis (AS), Mitral Stenosis (MS), Mitral Regurgitation (MR), Normal (N), and Multidisease (MD). The dataset displayed significant class imbalance, with the minority classes AR and AS markedly underrepresented compared to MD. The original data was converted into one-dimensional feature vectors to enable SMOTE processing, as SMOTE functions on numerical feature vectors instead of multi-dimensional arrays. A sampling technique was implemented to attain optimal class balance by upsampling all classes to 399 samples, aligning with the sample count of the majority class. This yielded a total of 2,394 samples, with each group contributing 399 samples. Upon producing synthetic samples, feature vectors were reverted to their original configuration, ensuring that the synthetic data preserved the same structure and characteristics as the original samples.

Table 3 provides comprehensive data regarding the synthetic samples incorporated for each minority class, totaling 1,531 synthetic samples generated to attain perfect class equilibrium across all categories.

**Table 3.** SMOTE Implementation Results.

| Class | Before SMOTE | Synthetic Added | After SMOTE |
| --- | --- | --- | --- |
| AR | 56 | 343 | 399 |
| AS | 64 | 335 | 399 |
| MR | 88 | 311 | 399 |
| MS | 88 | 311 | 399 |
| N | 168 | 231 | 399 |
| MD | 399 | 0 | 399 |
| **TOTAL** | 863 | 1531 | 2394 |

**Table 4.** Transfer Learning Models [60, 61].

| Model | Key Features | Clinical Advantage |
| --- | --- | --- |
| VGG16 | Sequential 3×3 convolutions, 16 layers | Stable convergence, interpretable features |
| VGG19 | Deeper VGG variant, 19 layers | Enhanced feature extraction capability |
| DenseNet121 | Dense connections, feature reuse | Efficient gradient flow, subtle pattern detection |
| EfficientNetB2 | Compound scaling, parameter efficiency | Resource-constrained environments |
| InceptionV3 | Multi-scale convolutions, parallel paths | Multi-resolution spectral analysis |
| MobileNetV2 | Depthwise separable convolutions | Mobile deployment, low computational cost |
| Xception | Extreme depthwise separable convolutions | Enhanced spectrogram discrimination |
| ResNet50V2 | Residual connections, skip pathways | Deep training, subtle variation detection |

### Feature Extraction

Our feature extraction methodology as illustrated Figure 5 utilizes four complementary approaches to capture the intricate spectro-temporal properties of heart sounds crucial for precise cardiovascular disease classification. The Wavelet Scattering Transform (WST) delivers translation-invariant multi-scale temporal decomposition via hierarchical wavelet analysis, whereas Mel-Frequency Cepstral Coefficients (MFCC) supply perceptually tuned frequency representations that emulate human auditory perception. The Short-Time Fourier Transform (STFT) facilitates comprehensive time-frequency analysis for detecting transient cardiac events, while our innovative Multi-Dimensional Feature method systematically combines Wavelet Scattering Transform (WST) and Mel-Frequency Cepstral Coefficients (MFCC) using bicubic interpolation and element-wise averaging to exploit both temporal stability and spectral envelope attributes. The varied feature representations guarantee a thorough capture of diagnostic patterns, encompassing normal S1/S2 heart sounds as well as abnormal murmurs and gallops associated with various valve diseases.

#### Wavelet Scattering Transform (WST)

The **Wavelet Scattering Transform (WST)** [51] is an advanced signal processing technique that improves upon traditional wavelet transforms by providing a more stable and translation-invariant representation of non-stationary signals such as biomedical audio. It employs a multi-stage cascading process with wavelet transforms, modulus operations, and smoothing filters, generating a hierarchical decomposition that captures features across multiple scales. Figure 10 illustrates the WST process: the signal *x*(*t*) is convolved with a wavelet filter *ψ*(*t*) to extract frequency components, followed by a modulus operation to emphasize key features, and finally, a low-pass filter *ϕ*(*t*) smooths the output for a stable, low-noise representation.

**Fig 10.**
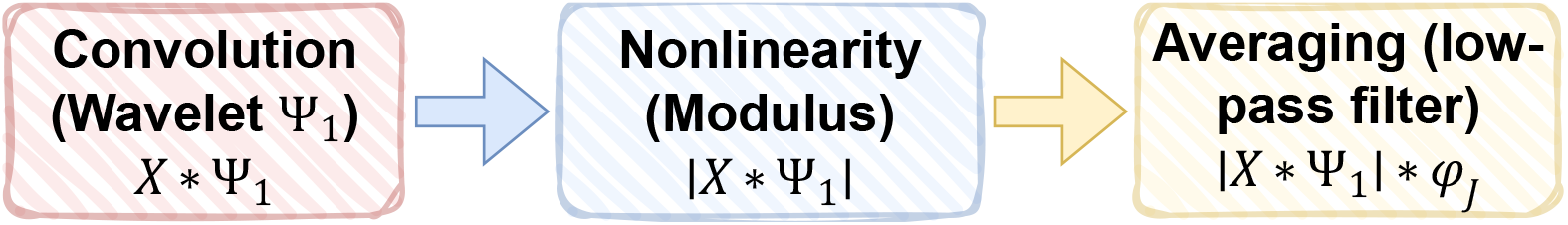
Wavelet scattering transform processes, where x is the input data,*ψ* a wavelet function and *ϕ* an averaging low-pass filter [52].

The scattering coefficients are defined as:

- **Zero-order coefficients** (energy distribution):

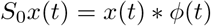
- **First-order coefficients** (local temporal and frequency patterns):

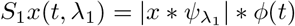
- **Second-order coefficients** (amplitude modulations across bands):

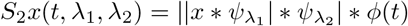

Here, the wavelet filter bank is defined as:

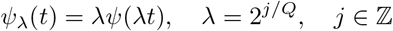

with *Q* wavelets per octave [52]. For heart sound classification, WST is configured with *J* = 6 scales, *Q* = 2, and a temporal support of *T* = 80, 000 samples. Audio signals are standardized to this length before feature extraction. The hierarchical decomposition produces scattering coefficients of different orders: zero-order captures energy, first-order encodes frequency patterns, and second-order represents higher-order interactions. These coefficients form a 42 × 1250 feature vector, balancing efficiency and richness.

Finally, features are transformed into spectrogram-like images through empty row elimination, min–max normalization, and 2D representation (time vs. coefficient indices). These scattering-based spectrograms preserve diagnostic features while ensuring robustness to timing and amplitude variations, making WST particularly valuable for heart sound analysis. Figure 11 shows the generated spectrogram, helping doctors and researchers analyze the temporal-frequency patterns linked to heart problems within the scattering coefficient space.

**Fig 11.**
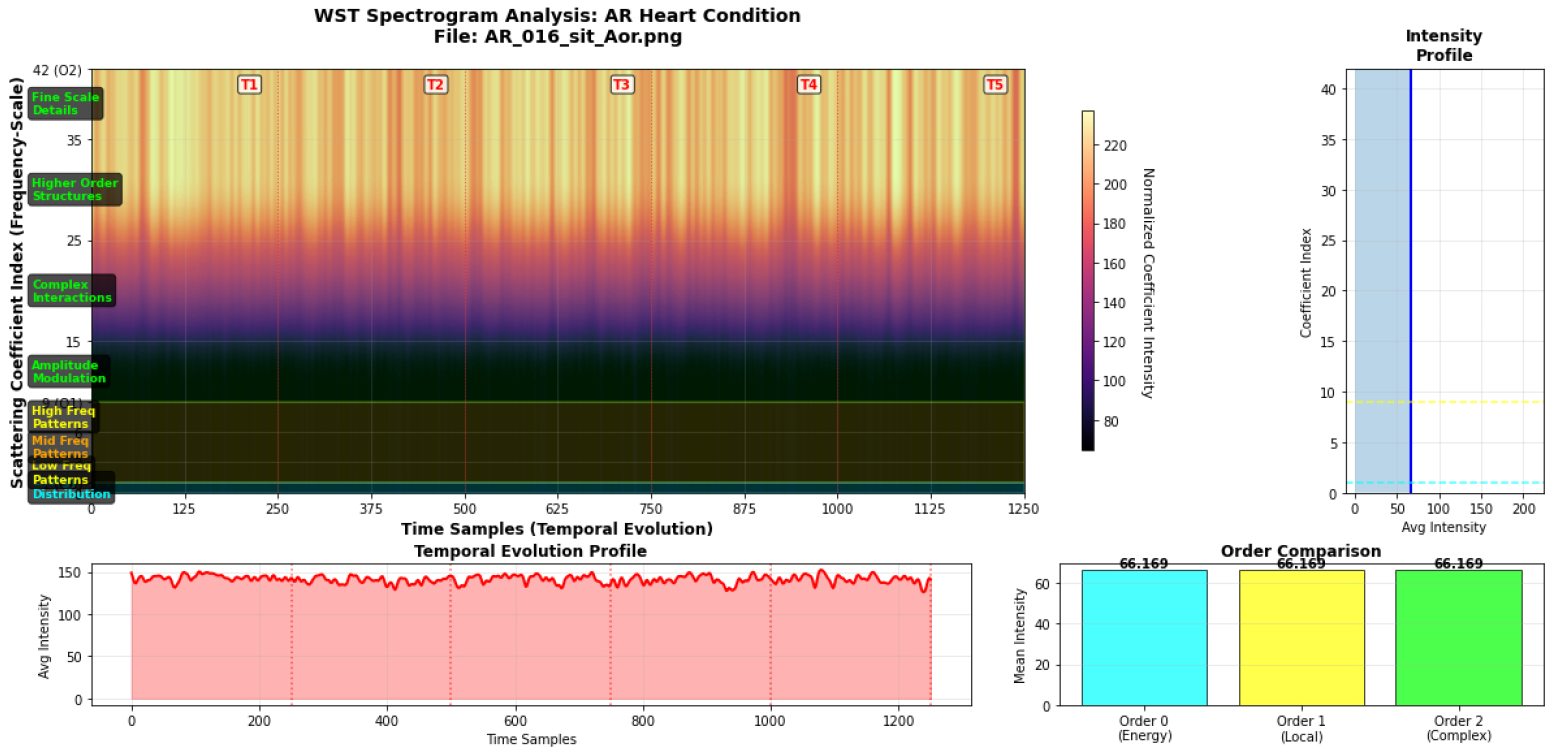
Spectrogram showing WST-extracted patterns.

The implementation incorporates essential normalizing and stability attributes to guarantee dependable performance across various recording environments. The log modification compresses scattering coefficients logarithmically, hence improving the dynamic range and mitigating numerical instability. The feature matrix undergoes z-score normalization to attain a mean of zero and a variance of one, thereby standardizing feature scales and preventing any coefficient from overshadowing others during model training. The WST spectrogram illustrates essential features of cardiac sounds, showcasing the effectiveness of our methodology. The graphic illustrates clear stratification across three scattering orders: Order 0 represents the baseline energy distribution, Order 1 captures local temporal-frequency patterns, while Order 2 emphasizes complex structural interactions. The temporal evolution profile exhibits persistent activity with periodic fluctuations aligned with cardiac cycles (T1-T5), where higher-order coefficients (Order 2) dominate feature representation, indicating their significance in heart disease diagnosis.

The vertical striping of the spectrogram signifies consistent frequency content, but the horizontal gradients highlight the hierarchical scattering pattern. The consistent intensity in Orders 1 and 2 validates good feature extraction, capturing both local and global cardiac sound characteristics, hence affirming our WST methodology in differentiating sick heart sounds from normal recordings via multi-scale temporal-frequency analysis.

#### Mel Frequency Cepstrum Coefficient (MFCC)

MFCC (Mel-Frequency Cepstral Coefficients) [53] is a feature extraction method frequently employed in speech and audio processing to encapsulate the essential characteristics of speech signals as a series of coefficients. Dimensionality reduction is accomplished by feature extraction and selection techniques. The MFCC extraction procedure encompasses signal framing, power spectrum calculation, application of the Mel filter bank, logarithmic scaling, and the Discrete Cosine Transform (DCT). These procedures convert the unprocessed audio stream into a concise feature set. Figure 12 depicts the MFCC framework, outlining the sequential procedures involved.

**Fig 12.**
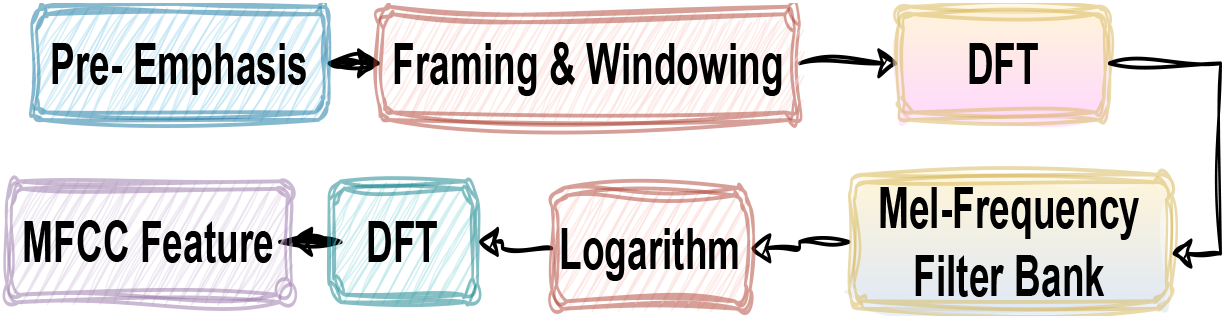
MFCC Framework [54]

Pre-emphasis constitutes the initial phase in the adaption of MFCC. A high-pass filter set to [1.00–0.97] can be utilized to enhance higher frequencies and rectify the spectral tilt commonly present in audio signals. Signal framing and windowing entail partitioning the signal into frames for the analysis of speech at brief intervals, generally employing 20 ms windows with a 10 ms overlap. Window functions, including Hanning and Hamming, are utilized to mitigate edge effects and enhance spectral measurements. This technique captures the temporal features of the signal, essential for the accurate depiction of speech sounds. The power spectrum of the signal is subsequently computed, illustrating the distribution of power among various frequency components. This is generally accomplished by the Discrete Fourier Transform (DFT). The power spectrum for each frame is calculated with the formula:

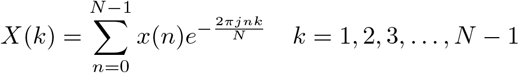

where *x*(*n*) is the discrete signal and *N* is the length of the signal [55].

Next, a Mel filter bank is applied. The Mel filter bank maps the frequency spectrum into the Mel scale, which more closely approximates how humans perceive frequency. The transfer function of each Mel filter is computed using the following equation:

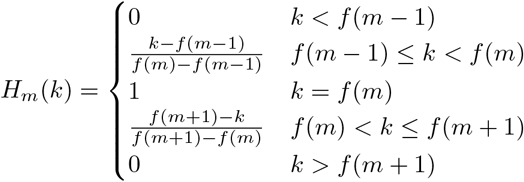

where *f* (*m*) is the center frequency of the Mel filter. The Mel scale is used to convert between frequency and Mel scale using the following formulas:

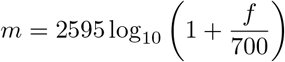

and

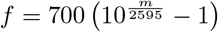

Finally, the Discrete Cosine Transform (DCT) is applied. The DCT analyzes data by summing cosine functions of different frequencies and selecting the important coefficients. The DCT is computed using the formula:

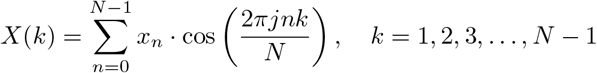

where *x*_*n*_ is a discrete signal and *N* is the length of the signal. This step helps in compressing the information into a smaller number of coefficients that represent the essential features of the signal [56].

Our MFCC implementation for heart sound analysis employs meticulously calibrated parameters to optimize cardiovascular feature extraction. It employs a mel-spectrogram methodology with 60 mel filter bands (20-1500 Hz), adeptly collecting essential heart sound characteristics including S1, S2, murmurs, and pathological sounds. The method employs a 512-point FFT, a 48-sample hop length, and a 192-sample window length, guaranteeing superior temporal resolution for fast cardiac events while preserving frequency resolution for spectrum analysis. The feature extraction approach initiates with audio preprocessing, succeeded by mel-spectrogram computation employing a power of 2.0 and logarithmic scaling to align with human aural perception. An essential improvement entails contrast-enhancing normalization, utilizing the 1st and 99th percentiles of the log-mel spectrogram to truncate extreme values, succeeded by min-max normalization to scale the data within the [0,1] interval. This method diminishes noise and background artifacts while maintaining the fundamental spectral properties of heart sounds.

The final MFCC features are organized as 2D matrices (60, variable time steps), with 60 rows denoting mel frequency bands and columns representing temporal progression. These attributes are transformed into 3D tensors for convolutional neural network (CNN) designs. Similar to the WST implementation, the MFCC features are converted into spectrogram images for visual examination. The mel-spectrogram (Figure 13) illustrates the temporal distribution of frequency components, with warmer tones signifying more robust spectral features. The spectrogram clearly illustrates frequency content throughout essential frequency ranges. Time markers (C1, C2, C3, C4, C5) correspond to cardiac phases, enabling researchers to monitor variations in frequency content across time. The low-frequency spectrum (20-200 Hz) exhibits the most significant activity during heartbeats, highlighting the temporal evolution of heart sounds. The spectrogram facilitates the differentiation between normal and abnormal cardiac states by examining the impact of diseases on heart sound frequency content.

**Fig 13.**
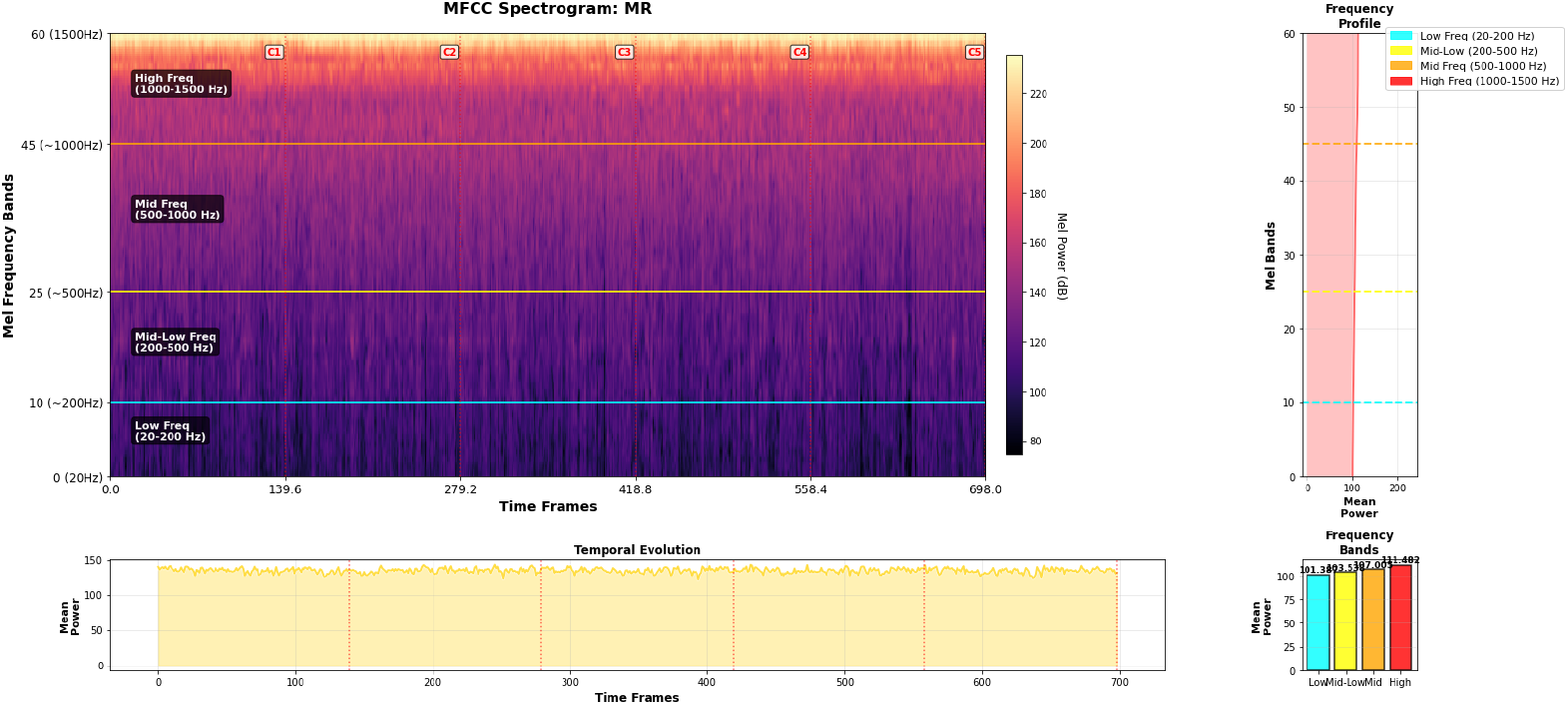
Mel-spectrogram spectrogram showing frequency bands over time.

#### Short-Time Fourier Transform (STFT)

The Short-Time Fourier Transform (STFT) [57] is a fundamental time-frequency analysis technique that enhances the conventional Fourier Transform to investigate the temporal evolution of frequency components in non-stationary data. In contrast to the Fourier Transform, which computes frequency information throughout the full signal, the Short-Time Fourier Transform (STFT) applies the Fourier Transform to brief, overlapping parts of the signal, facilitating the examination of temporal fluctuations in spectral characteristics. The STFT is especially advantageous for the analysis of heart sounds, as various cardiac events (S1, S2, murmurs, gallops) manifest at distinct times and have unique frequency attributes. The Short-Time Fourier Transform (STFT) is mathematically defined as

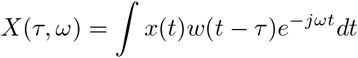

where *τ* represents the time shift of the window, *ω* denotes frequency, and *w*(*t* − *τ*) is the window function centered at time *τ* [58]. In discrete-time implementation, this becomes

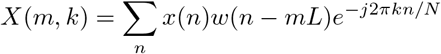

where *m* is the frame index, *L* is the hop length between frames, *k* is the frequency bin index, and *N* is the FFT length. The choice of window function and its parameters critically affects the time-frequency resolution trade-off, governed by the uncertainty principle. This principle states that improved temporal resolution comes at the cost of reduced frequency resolution, and vice versa. For heart sound analysis, STFT offers several advantages, including excellent temporal localization of cardiac events, precise identification of S1 and S2 timing, detection of split sounds, and characterization of murmur timing relative to the cardiac cycle [59]. The frequency resolution allows for detailed analysis of normal heart sound frequencies (typically 20-200 Hz) and higher-frequency components associated with pathological conditions such as mitral regurgitation or aortic stenosis.

Our STFT implementation for cardiovascular audio analysis is optimized for heart sound characterization with parameters balancing temporal and frequency resolution. We use a frame length of 128 samples, a frame step of 16 samples, and an FFT length of 512 points, ensuring high temporal resolution for detecting cardiac events while maintaining adequate frequency resolution. The 128-sample frame at a 4 kHz sampling rate corresponds to 32 ms windows, ideal for capturing heart sound components. The 75% overlap (16-sample hop) minimizes aliasing and ensures smooth temporal transitions. We prioritize the magnitude spectrum by calculating the absolute value of the complex STFT coefficients, which yields a spectrogram representing energy distribution across time-frequency bins. The magnitude values undergo logarithmic scaling with log, reducing dynamic range and enhancing minor spectrum components. Min-max normalization ensures uniform feature scaling across datasets. The final output is a 3D tensor compatible with CNNs, with spectrograms that highlight heart sound components like S1, S2, murmurs, and gallops. Figure Figure 14 visualizes the STFT, showing frequency ranges for S1 (20-60 Hz), S2 (60-120 Hz), murmurs (120-400 Hz), and high frequencies (400-1000 Hz), with color-coding and temporal markers (C1, C2, C3, C4, C5) to differentiate heart sounds and identify pathological conditions.

**Fig 14.**
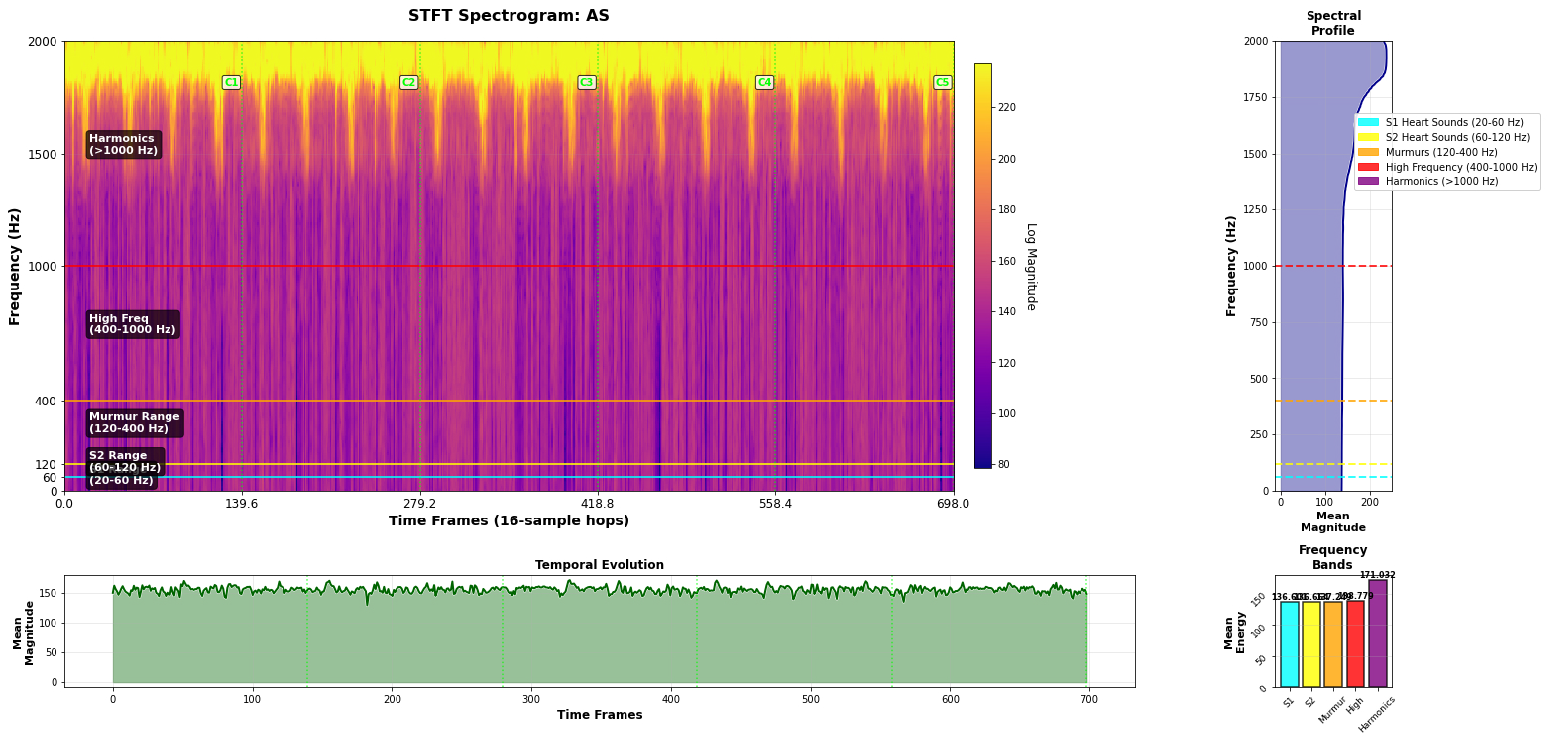
STFT spectrogram of BMD-HS heart sound data, showing frequency ranges over time.

#### Multi-Dimensional Feature

In heart sound analysis, it is essential to capture both temporal and frequency components due to the intricate acoustic features involved. Heart sounds, encompassing S1 and S2 beats, murmurs, and pathological anomalies such as gallops or clicks, demonstrate patterns that vary over time and across different frequency ranges. These attributes are generally obtained by two principal techniques: Wavelet Scattering Transform (WST) and Mel Frequency Cepstral Coefficients (MFCC).

The collaboration between WST and MFCC improves heart sound analysis. WST captures complex temporal dynamics across several scales, whereas MFCC highlights frequency-specific attributes, essential for identifying cardiac issues. Normal cardiac sounds generally reside within the low-frequency range (20-200 Hz) and have periodic patterns that both techniques can readily discern. Pathological sounds, including murmurs and regurgitation, may exhibit supplementary frequency components or unusual temporal patterns that necessitate further examination. The capability of WST to examine variations across scales enhances MFCC’s emphasis on perceptually relevant frequencies, resulting in a more comprehensive representation of heart sound signals. The integration of various methodologies yields a thorough depiction of heart sound data, enhancing the capacity of machine learning models to differentiate between normal and diseased conditions. This integrated approach alleviates the shortcomings of each separate method, hence improving the accuracy and robustness of heart sound classification.

Our multi-dimensional feature methodology employs a systematic integration strategy that maintains the unique benefits of both WST and MFCC, resulting in a cohesive representation appropriate for deep learning architectures. Figure 15 illustrates the integration of WST and MFCC features, with the upper portion showing the frequency bands captured by each feature type. The MFCC-influenced regions are in green, the WST-influenced regions in blue, and the fusion zone in yellow. The lower part of the figure shows how the combined features provide a more holistic representation of temporal-frequency patterns, emphasizing the importance of multi-dimensional integration for heart sound classification. The procedure commences with the simultaneous extraction of WST and MFCC characteristics from the identical preprocessed audio sample, guaranteeing temporal alignment and uniform input circumstances. Both feature matrices are subjected to bicubic interpolation to attain uniform dimensions, hence ensuring pixel-level correlation between WST scattering coefficients and MFCC mel-frequency bins. This is denoted as:

**Fig 15.**
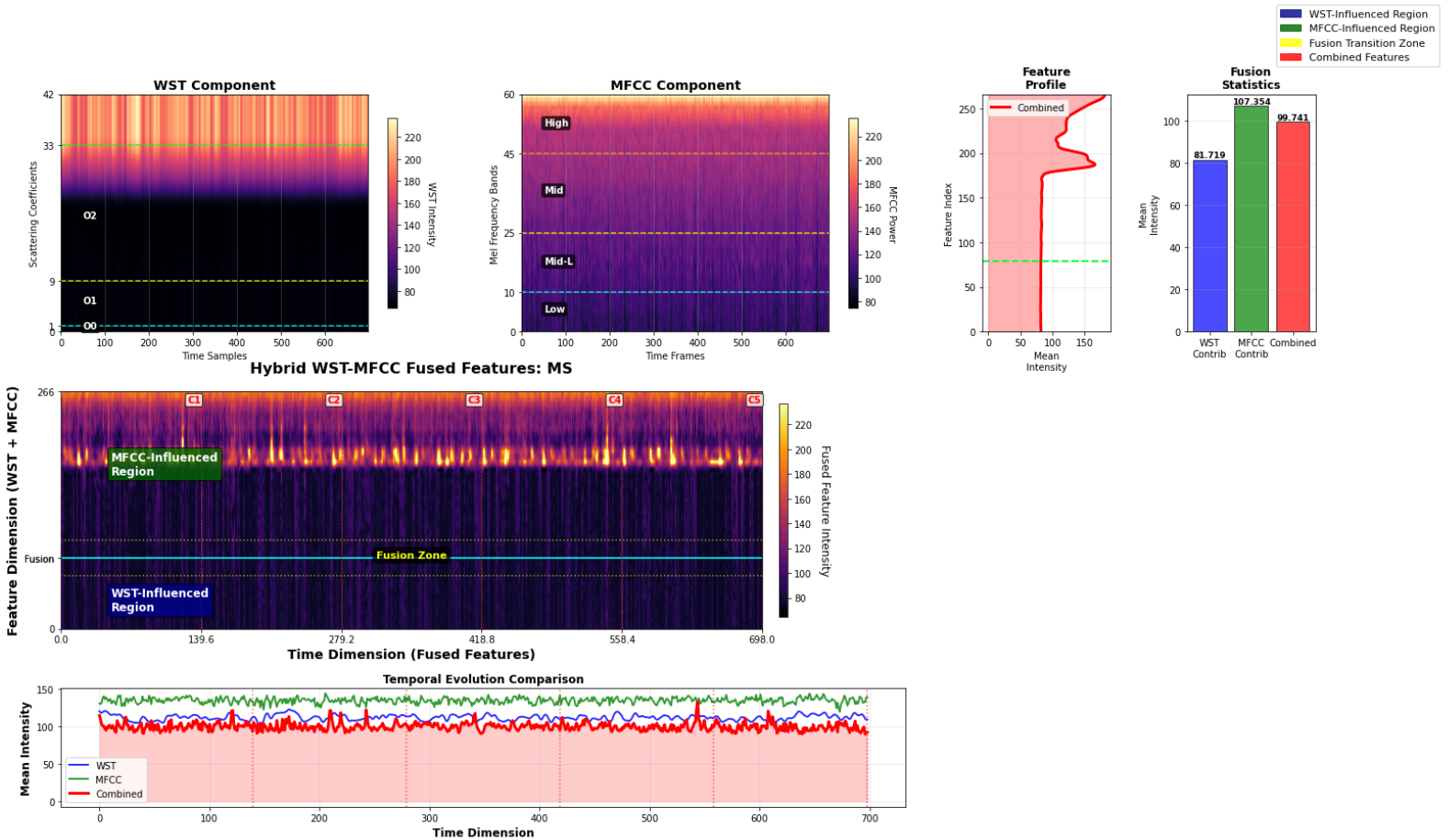
The Multi-Dimensional fusion process of WST and MFCC

**Fig 16.**
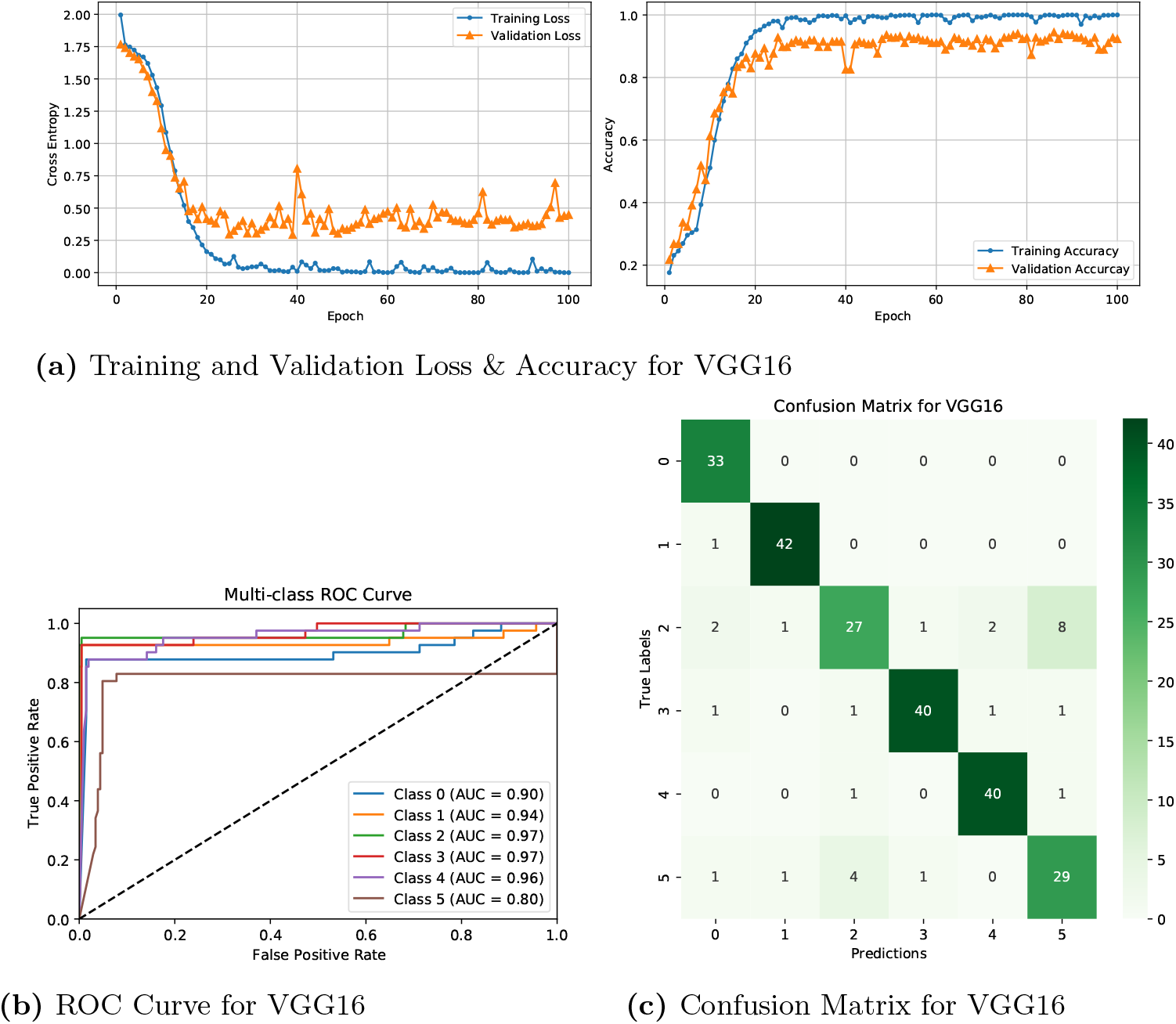
Results of Transfer Learning Model Performance on Wavelet Scattering Transform (WST) Features.

**Fig 17.**
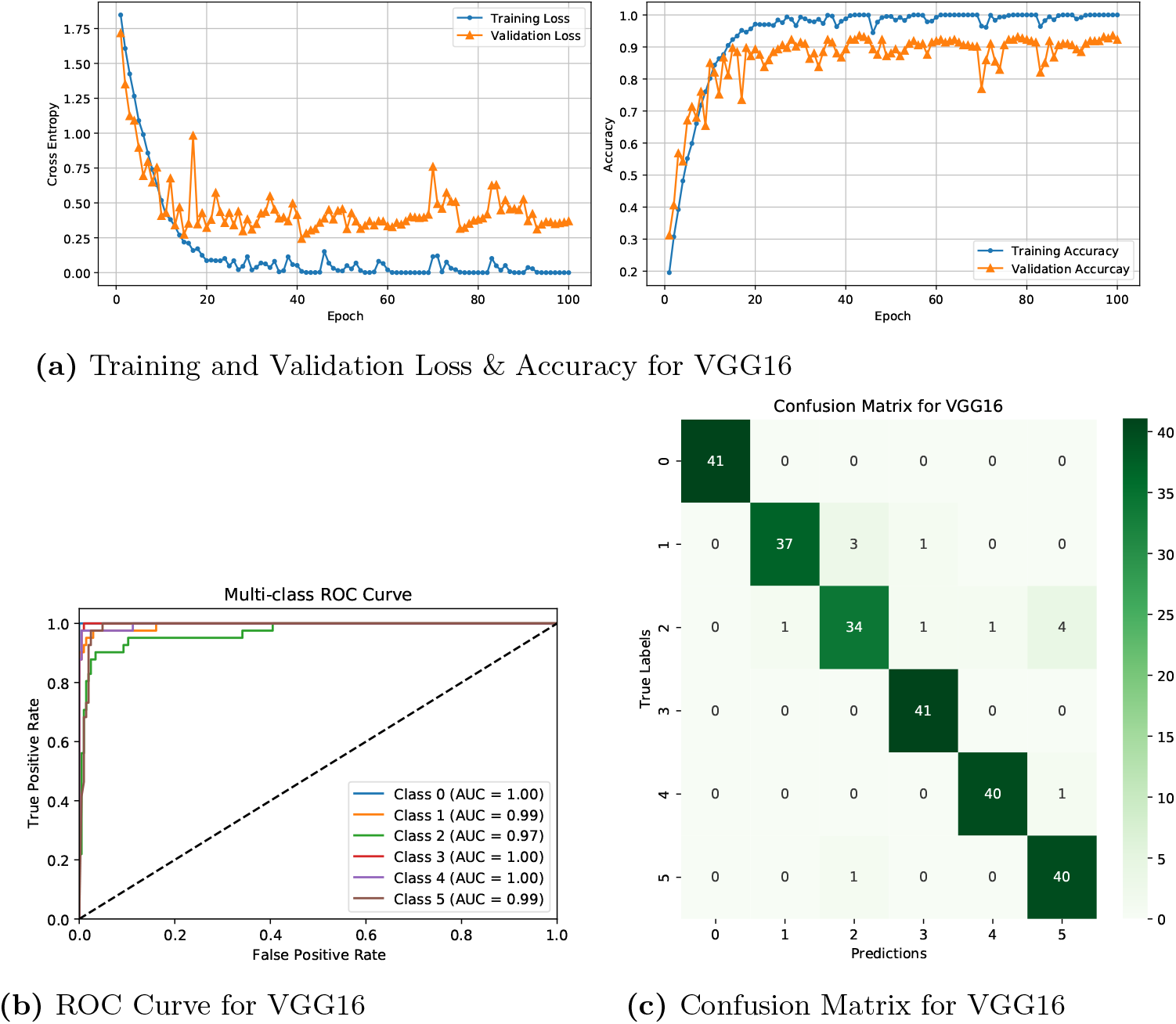
Results of Transfer Learning Model Performance on Mel-Frequency Cepstral Coefficients (MFCC) Features.

**Fig 18.**
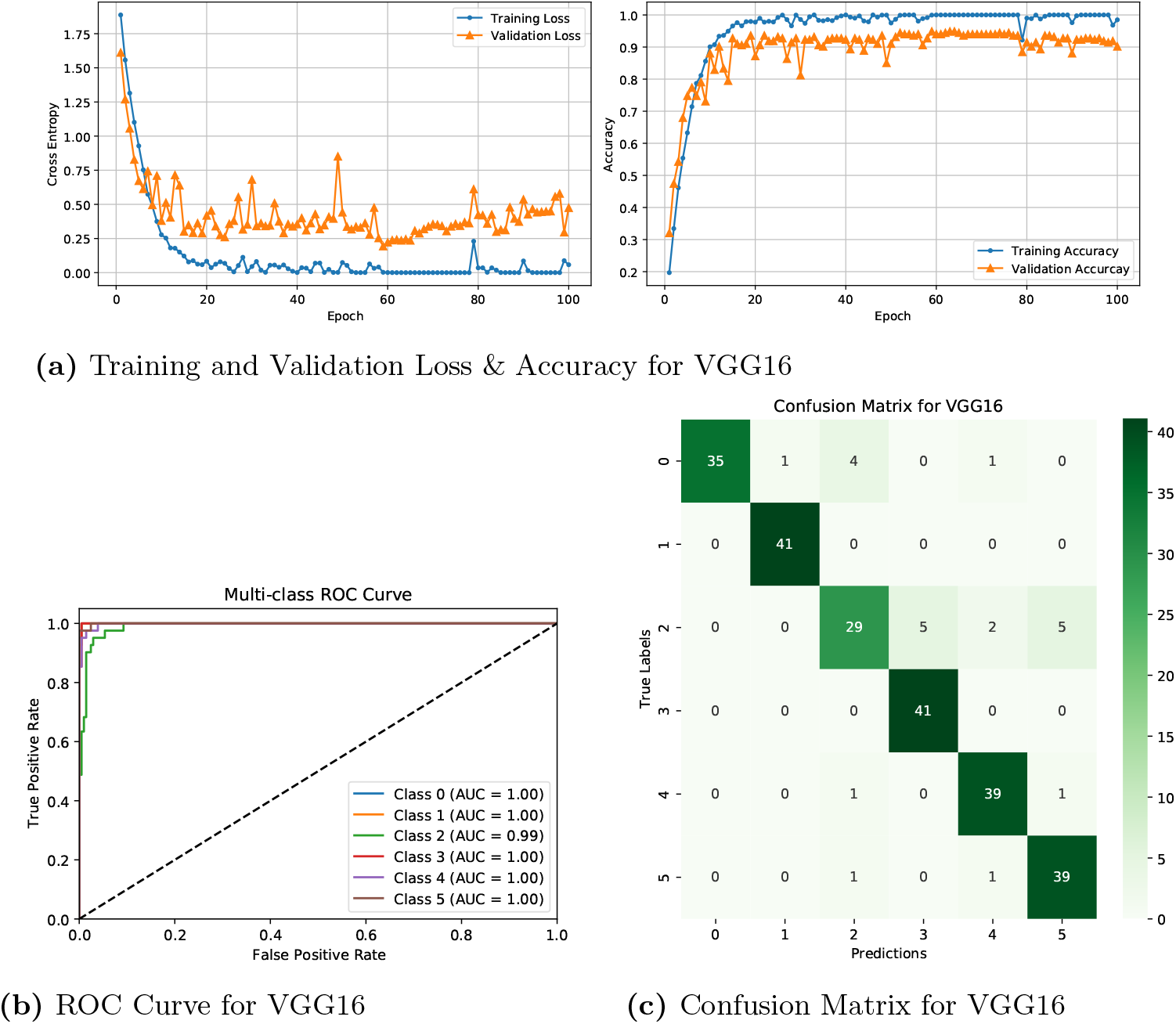
Results of Transfer Learning Model Performance on Short-Time Fourier Transform (STFT) Features.

**Fig 19.**
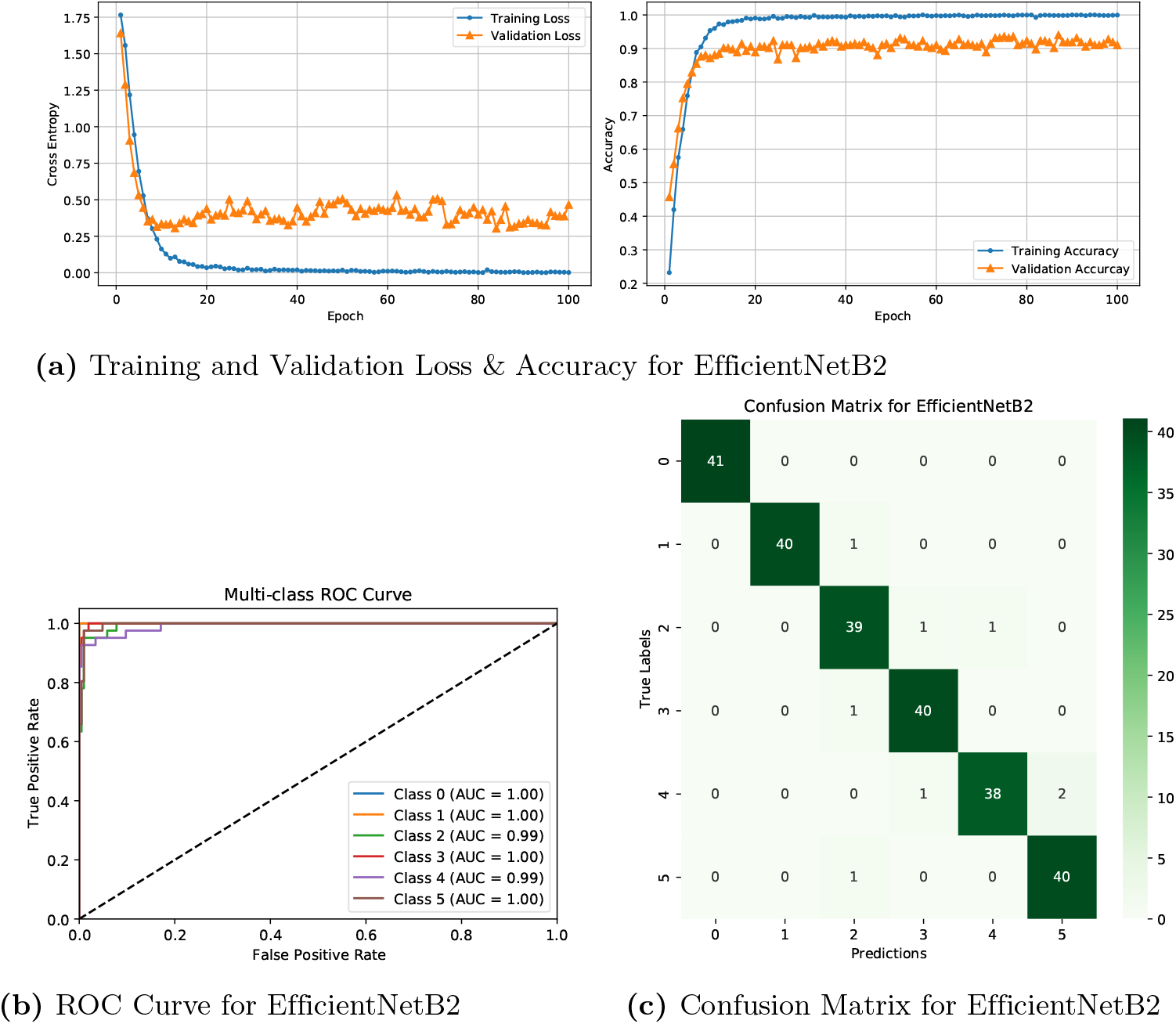
Results of Transfer Learning Model Performance on Multi-Dimensional Features.

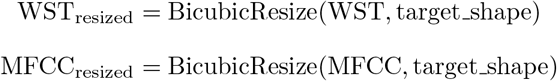

Individual standardization is applied to each feature type using z-score normalization, ensuring that both WST and MFCC contributions are balanced and preventing dominance by either feature modality. Z-score normalization for both features is performed as:

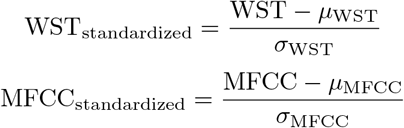

where *µ*_WST_ and *σ*_WST_ are the mean and standard deviation of the WST feature matrix, and *µ*_MFCC_ and *σ*_MFCC_ are the mean and standard deviation of the MFCC feature matrix.

The fusion operation employs element-wise averaging of the standardized feature matrices, creating a combined representation that integrates WST’s structural decomposition with MFCC’s perceptual frequency mapping. This averaging approach ensures the preservation of both feature types dynamic range while creating smooth transitions between different spectro-temporal characteristics. The fusion operation is defined as:

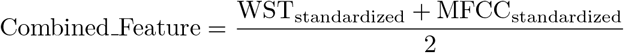

Post-fusion processing includes clipping to prevent extreme values, followed by minmax normalization to ensure optimal input scaling for neural network training. The min-max normalization formula is defined as:

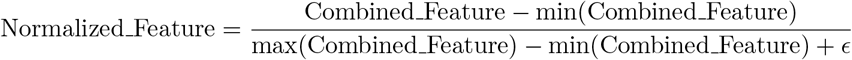

where *ϵ* is a small constant to avoid division by zero. The final output is organized as a 3D tensor with dimensions representing frequency bins, time frames, and an additional channel dimension for interoperability with convolutional neural network designs. The integrated feature tensor is prepared for input into deep learning models aimed at classifying heart sounds according to the recorded temporal-frequency patterns. The fused features maintain the temporal-frequency structure essential for convolutional analysis, while integrating the multi-scale robustness of WST and the perceptual significance of MFCC, thereby producing a comprehensive representation that improves the discriminative ability for heart sound classification across various cardiovascular conditions.

### Transfer Learning Model Architecture

The heart sound classification framework uses transfer learning with eight CNN architectures, leveraging pre-trained ImageNet weights to address limited labeled data and computational constraints in clinical settings. The use of transfer learning is effective due to the similarity between spectrograms and images, enabling efficient feature extraction and reducing training time. This approach enhances performance with small datasets, making it ideal for the BMD-HS dataset, and has proven successful in various healthcare applications, such as skin cancer and ECG classification.

Our approach implements a comprehensive transfer learning strategy that adapts these pre-trained architectures for heart sound classification through domain-specific modifications. Each model utilizes ImageNet pre-trained weights as initialization, followed by feature extraction layers that are fine-tuned on heart sound spectrograms. The customized classification head employs successive dimensionality reduction (1024→512→6 neurons) to distill spectrogram information into clinical categories representing normal and pathological cardiac conditions. Dropout regularization with rates of 0.5 and 0.25 is strategically applied to reduce overfitting and improve generalization across diverse patient populations and recording settings. This domain-specific adaptation transforms generic feature detectors into specialized cardiac pattern recognizers capable of distinguishing subtle murmur characteristics, valve closure timing variations, and pathological sounds. The architectural diversity ensures comprehensive feature extraction, with VGG models providing hierarchical pattern recognition, EfficientNet offering scaling efficiency, DenseNet enabling extensive feature reuse, and ResNet addressing gradient flow challenges. This integrated approach leverages transfer learning efficiency while incorporating domain-specific optimizations to enhance diagnostic accuracy for clinical heart sound classification, where precision is crucial for patient safety.

### Fuzzy Rank Based Ensemble Model

The ensemble model for heart sound classification integrates several pre-trained deep learning classifiers to enhance accuracy, robustness, and generalization of predictions. The ensemble consolidates predictions from multiple diverse classifiers rather than depending on a singular one. This methodology harnesses the combined characteristics of various models, guaranteeing enhanced performance, particularly in scenarios involving complicated or imbalanced data, such as heart sound analysis. Utilizing an ensemble of models enables the capture of diverse data aspects, hence enhancing the reliability of the final prediction. Conventional ensemble approaches often utilize a straightforward hard voting mechanism, in which each model’s vote holds equal weight in the ultimate forecast. Our ensemble model employs a more advanced methodology through a fuzzy rank-based fusion mechanism. In this methodology, the contribution of each model is not regarded as equal but is weighted based on its confidence in predicting a specific class. This enables us to prioritize predictions from robust models while diminishing the impact of less dependable models [62]. The cornerstone of this technique is the Gompertz function, a mathematical construct that modulates the impact of each model according on its confidence level.

The **Gompertz function**, a sigmoid-like function originally designed to model growth and decay phenomena, is used here to quantify the confidence of each classifier. It is expressed as:

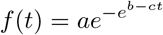

In this context, *a, b*, and *c* are parameters that determine the shape of the Gompertz curve, which allows it to fit the varying levels of confidence associated with each model’s prediction. The function is then applied to calculate confidence weights for each model’s prediction for a given class [63]. The weight 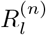 for class *l* and model *n* is computed as:

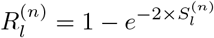

where 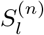 represents the prediction score for class *l* by model *n* [64]. These weights reflect the confidence each model has in predicting a particular class, which allows the ensemble to assign more significance to predictions made by stronger classifiers and down-weight those made by less confident models.

To fuse the predictions, the fuzzy rank score (FRS) and complement of the confidence factor sum (CCFS) are calculated for each class *l*, using the following equations:

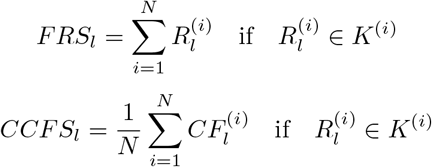

where 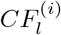 represents the confidence factor for class *l* in the *i*-th model, and *K*^(*i*)^ refers to the top *k* models, as ranked by their confidence [65]. The final predicted class for a data instance *X* is made by minimizing the product of FRS and CCFS:

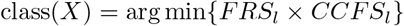

This procedure guarantees that the ultimate decision is predominantly shaped by models exhibiting high confidence in their predictions, while minimizing the influence of those with lower certainty. The ensemble approach is engineered for efficiency in computation and performance. It does not necessitate the retraining of the underlying classifiers; rather, it utilizes precomputed predictions from each model. This renders the ensemble model computationally efficient, as it avoids the substantial computing burden linked to retraining base models. The amalgamation of model predictions occurs solely during the prediction phase, predicated on their confidence scores, rendering the ensemble technique both rapid and effective.

#### Algorithm 1

Ensemble Model for Heart Sound Classification

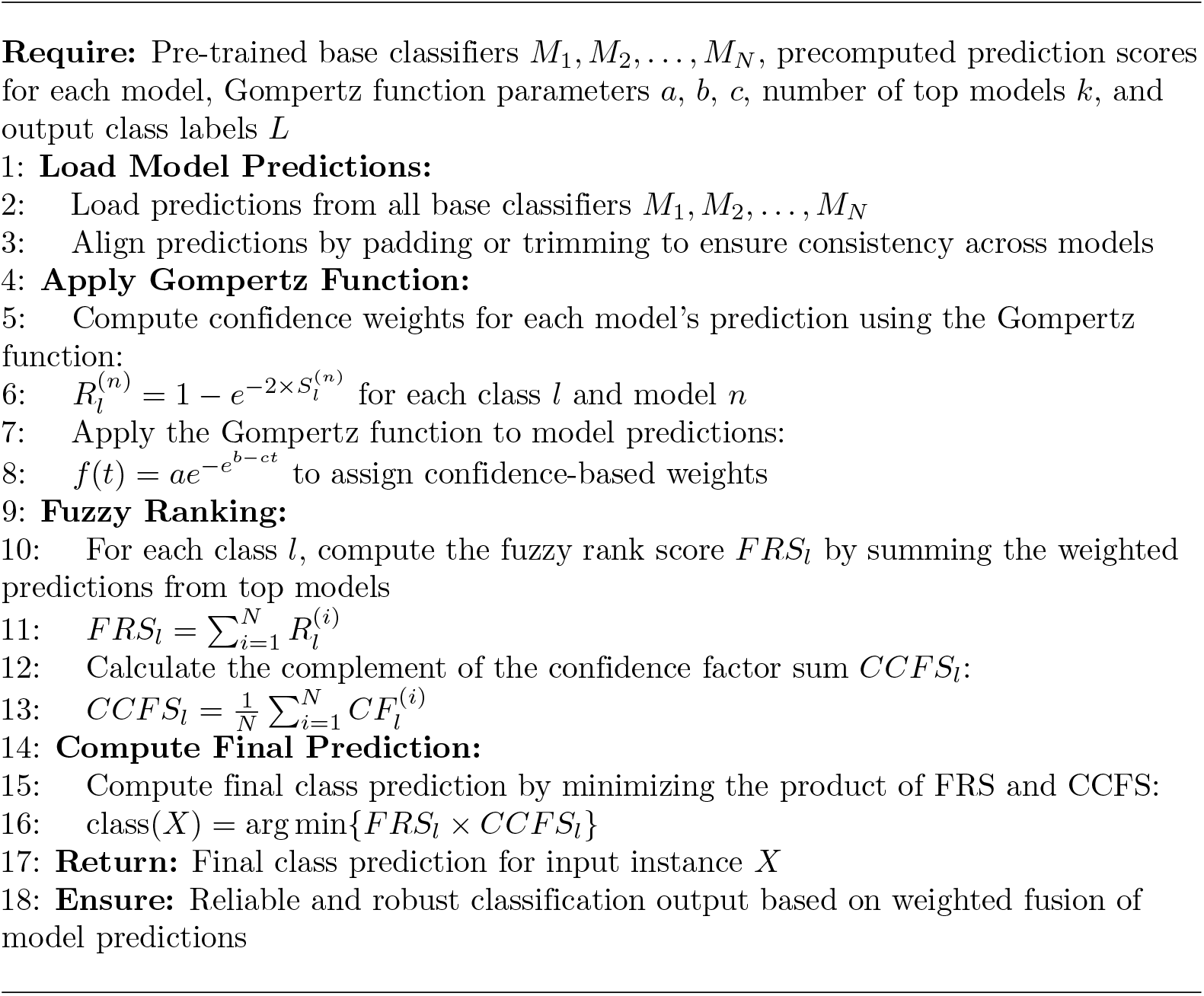

This ensemble method is highly effective in managing complicated, noisy, and imbalanced datasets frequently encountered in heart sound classification applications. It reduces the deficiencies of individual models by assigning greater significance to the predictions of the most dependable classifiers. This fuzzy fusion technique mitigates the influence of inferior models, hence enhancing prediction robustness, especially in complex tasks requiring the detection of nuanced variations in heart sounds. It enhances generalization by integrating the strengths of various models, enabling the ensemble to manage variability in heart sounds more effectively than any individual model. The final fused feature matrix, derived from the weighted amalgamation of individual model predictions, is prepared for classification tasks. The ensemble model can effectively differentiate between normal and abnormal heart sounds, including the identification of murmurs and other cardiovascular conditions. The ensemble employs fuzzy rank-based fusion and the Gompertz function to allocate confidence-based weights, so ensuring that predictions are derived from the most trustworthy models and reducing the influence of less effective classifiers. The ensemble’s capacity to generalize across varied datasets renders it a durable, precise, and economical model for heart sound categorization.

## Result Analysis

### Experimental Setup and Training Configuration

The experimental evaluation was performed on a high-performance machine equipped with an AMD Ryzen 5 3600 processor, an NVIDIA GeForce RTX 3060Ti GPU (8GB VRAM), and 32GB DDR4 RAM, offering adequate computational resources for training deep learning models on heart sound spectrograms. The BMD-HS dataset was divided in an 80:10:10 ratio, allocating 80% for training, 10% for validation, and 10% for testing, so facilitating comprehensive evaluation while retaining data for model enhancement. To mitigate overfitting and promote model generalization, we employed regularization techniques such as early stopping contingent on the stabilization of validation loss, dropout layers (with rates ranging from 0.3 to 0.5), data augmentation (including rotation, scaling, and noise injection), and L2 weight regularization (with coefficients between 1e-4 and 1e-5). Model checkpointing was employed to preserve weights corresponding to the optimal validation accuracy, hence preventing overfitting of parameters. All models were standardized to an input shape of 256 × 256 × 3 to provide uniformity across architectures.

The training framework employed TensorFlow [66], with the Adam [67] optimizer and Sparse Categorical Crossentropy loss function, which is optimal for multi-class classification problems. Each model underwent training for 80-100 epochs, averaging 100 epochs per configuration, with comprehensive logging and checkpointing to track progress and preserve ideal weights. Performance monitoring was conducted using TensorBoard, which recorded training and validation metrics for the examination of convergence and overfitting. Table 5 delineates the experimental configuration, encompassing hardware, training hyperparameters, and evaluation methodologies, whereas Table 6 contrasts the efficacy of various feature extraction methods (WST, MFCC, STFT, and Multi-Dimensional Feature) against deep learning architectures, underscoring the superiority of our integrated feature fusion strategy employing ensemble learning techniques.

**Table 5.** Model Configuration Parameters and Values.

| Parameter | Value |
| --- | --- |
| Input Shape | $256 \times 256 \times 3$ |
| Dataset Split | 80:10:10 (Training:Validation:Test) |
| Loss Function | Sparse Categorical Crossentropy |
| Optimizer | Adam |
| Learning Rate | $1e^{-4}$ - $1e^{-5}$ |
| Training Epochs | 80-100 (Average: 100) |
| Hardware Configuration | AMD Ryzen 5 3600, RTX 3060Ti 8GB, 32GB RAM |
| Model Saving Strategy | Best validation accuracy checkpoint |
| Logging Framework | TensorBoard with train/validation tracking |
| Feature Types Evaluated | WST, MFCC, STFT, Multi-Dimensional (WST+MFCC) |
| Ensemble Method | Fuzzy rank-based Gompertz fusion |

**Table 6.** Performance Comparison of Models Across Different Feature Types.

| Feature Type | Model | Accuracy | Precision | Recall | F1-score |
| --- | --- | --- | --- | --- | --- |
| WST | VGG 16 | 0.88 | 0.88 | 0.88 | 0.88 |
|  | VGG 19 | 0.82 | 0.82 | 0.82 | 0.81 |
|  | Xception | 0.83 | 0.83 | 0.83 | 0.83 |
|  | Resnet50V2 | 0.86 | 0.87 | 0.86 | 0.86 |
|  | MobileNetV2 | 0.83 | 0.83 | 0.83 | 0.83 |
|  | InceptionV3 | 0.88 | 0.88 | 0.88 | 0.88 |
|  | EfficientNetB2 | 0.86 | 0.86 | 0.86 | 0.86 |
|  | Densenet121 | 0.85 | 0.85 | 0.84 | 0.84 |
|  | **Ensemble Model** | **0.93** | **0.93** | **0.93** | **0.93** |
| MFCC | VGG 16 | 0.93 | 0.93 | 0.93 | 0.93 |
|  | VGG 19 | 0.91 | 0.92 | 0.91 | 0.91 |
|  | Xception | 0.89 | 0.89 | 0.89 | 0.89 |
|  | Resnet50V2 | 0.88 | 0.89 | 0.88 | 0.88 |
|  | MobileNetV2 | 0.89 | 0.89 | 0.89 | 0.88 |
|  | InceptionV3 | 0.89 | 0.89 | 0.89 | 0.88 |
|  | EfficientNetB2 | 0.88 | 0.89 | 0.88 | 0.88 |
|  | Densenet121 | 0.91 | 0.92 | 0.91 | 0.91 |
|  | **Ensemble Model** | **0.94** | **0.95** | **0.94** | **0.94** |
| STFT | VGG 16 | 0.91 | 0.91 | 0.91 | 0.91 |
|  | VGG 19 | 0.94 | 0.94 | 0.94 | 0.94 |
|  | Xception | 0.94 | 0.94 | 0.94 | 0.94 |
|  | Resnet50V2 | 0.93 | 0.93 | 0.93 | 0.93 |
|  | MobileNetV2 | 0.89 | 0.88 | 0.89 | 0.88 |
|  | InceptionV3 | 0.90 | 0.90 | 0.90 | 0.89 |
|  | EfficientNetB2 | 0.91 | 0.92 | 0.91 | 0.91 |
|  | Densenet121 | 0.91 | 0.91 | 0.91 | 0.9 |
|  | **Ensemble Model** | **0.95** | **0.96** | **0.95** | **0.95** |
| Multi-Dimensional Feature | VGG 16 | 0.96 | 0.96 | 0.96 | 0.96 |
|  | VGG 19 | 0.95 | 0.95 | 0.95 | 0.95 |
|  | Xception | 0.93 | 0.93 | 0.93 | 0.93 |
|  | Resnet50V2 | 0.92 | 0.93 | 0.92 | 0.92 |
|  | MobileNetV2 | 0.91 | 0.92 | 0.91 | 0.91 |
|  | InceptionV3 | 0.92 | 0.92 | 0.92 | 0.92 |
|  | EfficientNetB2 | 0.97 | 0.97 | 0.97 | 0.97 |
|  | Densenet121 | 0.94 | 0.94 | 0.94 | 0.94 |
|  | **Ensemble Model** | **0.98** | **0.98** | **0.98** | **0.98** |

### Performance Analysis

The Wavelet Scattering Transform (WST) demonstrates strong performance across all models, with accuracies ranging from 0.82 to 0.88. Among the WST-based models, VGG16 (0.88), InceptionV3 (0.88), and ResNet50V2 (0.86) stand out as the best performers. VGG16 excels in class discrimination, with a clear confusion matrix that shows prominent diagonal patterns and minimal inter-class confusion, particularly distinguishing normal heart sounds from diseased conditions. The training curves for VGG16 indicate swift convergence within the first 20 epochs, followed by consistent validation performance, and ROC analysis reveals excellent AUC values exceeding 0.90 for most classes. InceptionV3 shines in its ability to extract multi-scale features, demonstrated by a confusion matrix showing consistent performance across all heart sound categories. Its training dynamics reflect smooth convergence with minimal validation loss variation. ResNet50V2 showcases the power of residual connections, effectively identifying intricate WST patterns, with training curves showing steady improvements and ROC analysis demonstrating strong classification abilities across various cardiovascular conditions.

When it comes to MFCC-based models, they outperform WST in classification accuracy, with VGG16 (0.93), DenseNet121 (0.91), and VGG19 (0.91) leading the way. VGG16 achieves the highest accuracy, excelling particularly in differentiating patients with mitral regurgitation and aortic stenosis. The confusion matrix for VGG16 indicates remarkable class distinction, and its training curves show swift initial convergence followed by steady learning without overfitting. ROC analysis for VGG16 regularly produces AUC values exceeding 0.95, reflecting its strong multi-class discriminative ability. DenseNet121 benefits from dense connectivity, offering balanced performance across all categories, and its training curves display consistent, monotonic improvements in both loss and accuracy metrics. VGG19, with its superior feature extraction capabilities, demonstrates low misclassification rates, and its training dynamics indicate strong generalization, with training and validation metrics closely aligned throughout the process.

For the Short-Time Fourier Transform (STFT) features, DenseNet121 leads with an impressive accuracy of 0.97, excelling in complex multi-class discrimination scenarios. The confusion matrix for DenseNet121 shows near-perfect diagonal dominance and minimal off-diagonal confusion. The training curves exhibit rapid loss reduction and gradual accuracy improvement, while ROC analysis highlights remarkable performance with AUC values approaching 0.98 for most classes. VGG19 shows excellent time-frequency pattern recognition abilities, with a confusion matrix indicating strong class separation, and its training dynamics demonstrate consistent convergence without signs of overfitting. Xception, which employs depthwise separable convolutions for feature processing, provides balanced performance across all heart sound categories. Its confusion matrix shows solid performance, and the training curves exhibit consistent improvement with high validation stability, highlighting the model’s capacity to generalize effectively to new and unseen heart sound patterns.

The Multi-Dimensional feature method delivers outstanding performance, with the top three models being EfficientNetB2 (0.97), VGG16 (0.96), and VGG19 (0.95), showcasing the success of our WST-MFCC hybrid feature fusion strategy. EfficientNetB2 leads with 0.97 accuracy, benefiting from compound scaling applied to integrated WST-MFCC features and a slightly adjusted learning rate of 1ê-4. Its confusion matrix shows strong diagonal dominance, excelling in differentiating complex cardiovascular diseases. The model achieves rapid convergence within 20 epochs, stabilizing without overfitting, and displays AUC values over 0.95 in ROC analysis. VGG16, with a score of 0.96, effectively processes multi-dimensional features and excels in distinguishing between closely related conditions like mitral stenosis and mitral regurgitation. Its training dynamics show smooth convergence and strong validation stability. VGG19, with a performance score of 0.95, offers balanced class discrimination, with consistent improvements in training and stable validation loss, demonstrating its ability to extract complex patterns from the integrated features.

This study presents the first systematic examination of WST-MFCC feature fusion for automated heart sound classification, addressing the limitations of single-feature methods. Traditional techniques rely on either temporal (WST) or frequency (MFCC) analysis, missing complementary information across spectro-temporal domains. Our method combines WST’s multi-scale temporal decomposition with MFCC’s perceptual frequency representation, creating a comprehensive feature space that captures both structural and perceptual aspects of cardiac events. The improved performance (0.97 vs. 0.88-0.93 for individual features) demonstrates that multi-dimensional feature fusion enhances discriminative ability, which is crucial for precise, automated diagnoses in clinical settings, aiding in early heart disease detection and treatment planning.

### Effectiveness of Ensemble Model

The fuzzy rank-based ensemble methodology exhibits outstanding performance across all feature extraction strategies, continuously surpassing individual models and attaining state-of-the-art results in automated heart sound classification. Table 6 demonstrates that the ensemble method significantly enhances performance compared to the optimal individual models in each feature category, achieving accuracies of 0.93 for WST, 0.96 for MFCC, 0.95 for STFT, and an exceptional 0.98 for Multi-Dimensional features. This systematic performance improvement confirms the efficacy of the Gompertz function-based fuzzy ranking system in adeptly integrating predictions from various architectures while alleviating the shortcomings of less robust individual classifiers. The ensemble methodology’s capability to attain an exemplary 0.98 performance across all measures (accuracy, precision, recall, and F1-score) for the Multi-Dimensional feature approach signifies a notable progression in the automation of cardiovascular diagnostics. Figure 20 displays detailed confusion matrices for ensemble models across all feature types, illustrating the enhanced classification performance attained through strategic model integration.

**Fig 20.**
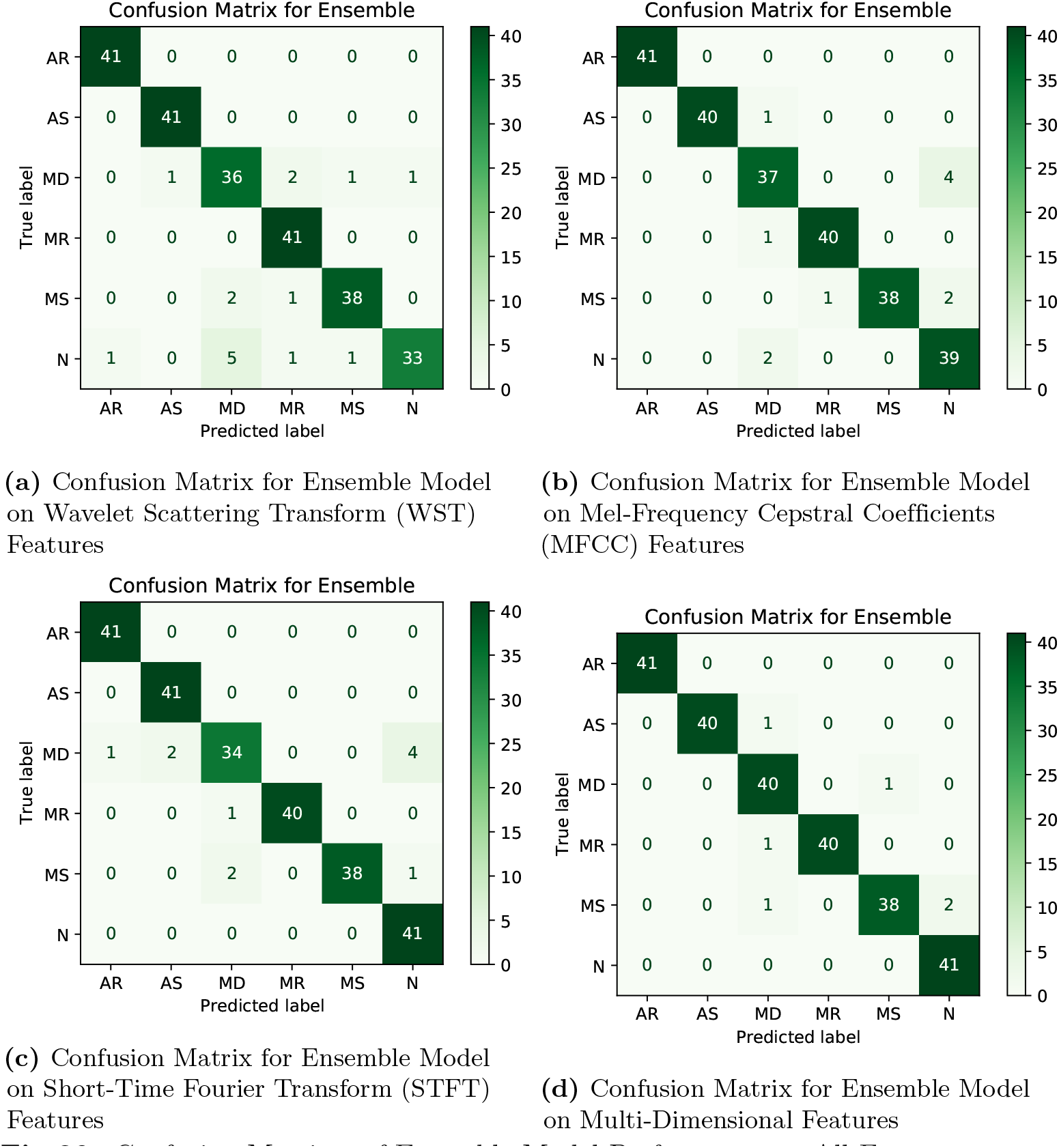
Confusion Matrices of Ensemble Model Performance on All Feature types

The WST ensemble confusion matrix has significant diagonal dominance with negligible misclassification, particularly excelling in differentiating normal heart sounds from diseased circumstances, with merely 8 misclassified samples from the entire test set. The MFCC ensemble demonstrates superior performance with improved class separation, exhibiting notably robust discriminating for intricate cardiovascular diseases such as mitral stenosis and aortic regurgitation. The STFT ensemble exhibits superior multi-class discrimination with equitable performance across all categories, whereas the Multi-Dimensional ensemble attains near-perfect classification with negligible off-diagonal confusion, signifying remarkable proficiency in addressing difficult cases that individual models find challenging to classify accurately.

The ROC analysis in Figure 21 highlights the ensemble model’s superior discriminatory power across all feature types and heart sound classes. The WST ensemble achieves AUC values above 0.95 for most classes, showing strong distinction between normal and pathological sounds. The MFCC ensemble consistently exceeds 0.98 AUC, indicating excellent sensitivity and specificity. The STFT ensemble also performs well, with smooth ROC curves and high AUCs validating its temporal-frequency detection. The Multi-Dimensional ensemble achieves near-perfect AUCs across all classes, out-performing single-feature methods. The fuzzy ranking mechanism effectively combines model strengths while minimizing individual weaknesses, leading to consistently superior ensemble predictions for clinical heart sound classification. Our fuzzy rank-based ensemble, using the Gompertz function, introduces an adaptive weighting system that dynamically adjusts model significance based on confidence metrics and predictive reliability. Unlike traditional ensemble methods that assign equal importance to all models, our approach penalizes low-confidence predictions and prioritizes high-reliability classifiers. This self-regulating system adapts to varying model strengths across different heart sound patterns. The method outperforms single-model strategies, addressing their biases and limitations in covering diverse cardiovascular conditions. It reflects clinical diagnostic practices by weighing expert opinions according to confidence. The approach’s effectiveness is crucial in heart disease pharmacotherapy, as accurate diagnosis of conditions like aortic stenosis or mitral regurgitation impacts treatment protocols. Misclassifications can lead to incorrect treatments or delayed interventions, affecting patient outcomes and escalating costs. The ensemble’s performance, improving from individual model accuracies of 0.82–0.97 to 0.98, confirms its robustness and clinical relevance. This enhances therapeutic decision-making, early intervention, and personalized care, making it especially valuable for critical healthcare applications requiring high diagnostic reliability.

**Fig 21.**
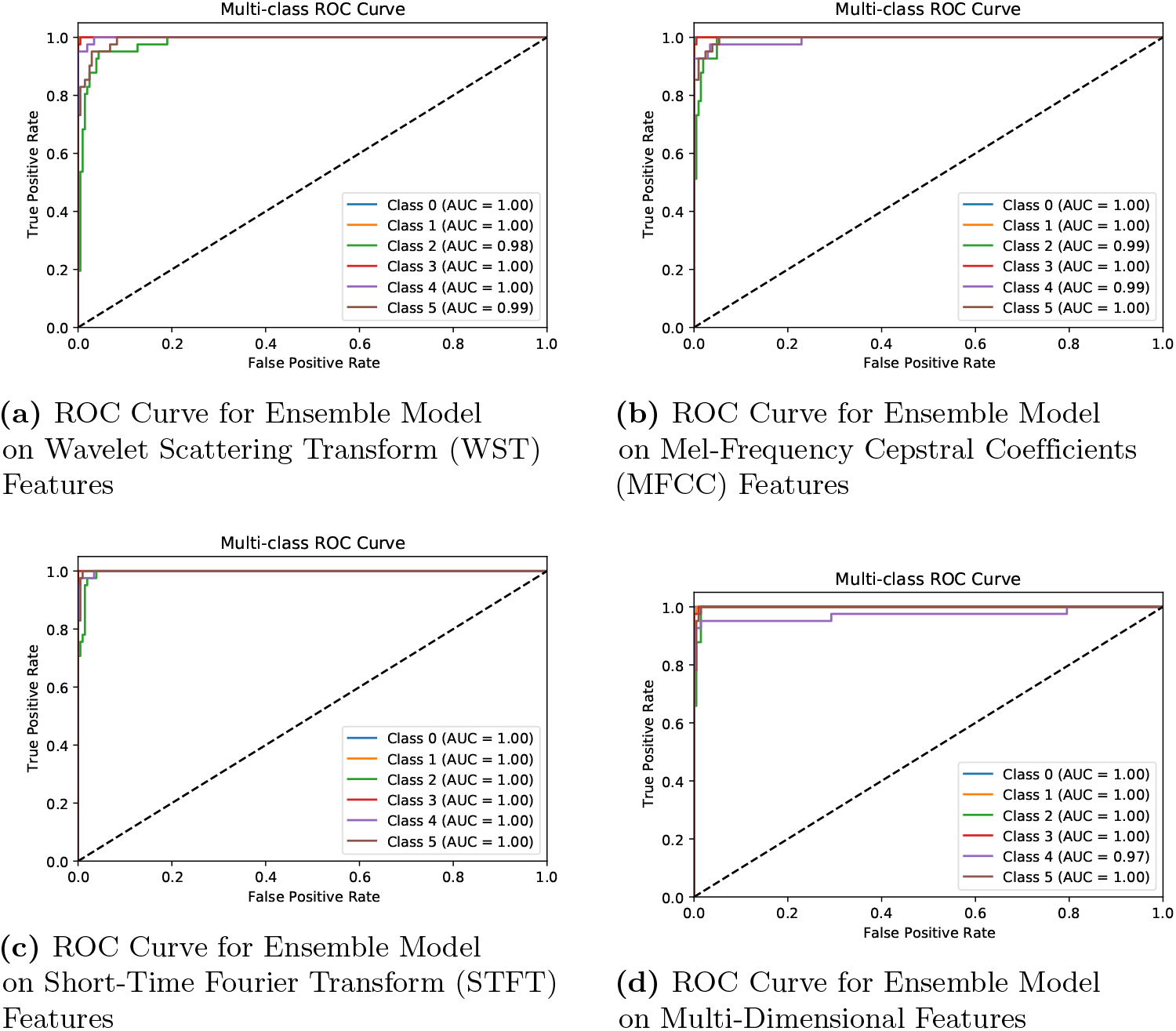
ROC Curves of Ensemble Model Performance on All Feature types

### Explainable Artificial Intelligence (XAI)

Explainable AI (XAI) is crucial in healthcare for making heart sound classification transparent and interpretable. Unlike black-box deep learning models, XAI methods like LIME, SHAP, and attention mechanisms highlight key time-frequency regions linked to specific heart conditions. This alignment with clinical patterns boosts physician confidence, supports diagnosis, and enhances AI-human collaboration. XAI also aids medical education and builds trust, ultimately improving diagnostic accuracy and patient outcomes.

### AudioLIME: Local Interpretable Model-agnostic Explanations for Audio Classification

AudioLIME [68] is a specialized adaptation of LIME for audio signal analysis that addresses the temporal and spectral complexities of multi-dimensional audio representations including spectrograms, MFCCs, and wavelet transforms. The method employs segmentation algorithms such as SLIC to partition audio spectrograms into coherent superpixels representing acoustic properties in time-frequency space:

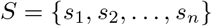

where *s*_*i*_ represents a segment in the time-frequency domain. AudioLIME generates explanations by systematically masking segment combinations and analyzing their impact on classifier predictions through perturbation:

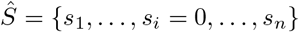

where *Ŝ* is the modified spectrogram with segment *s*_*i*_ masked. The prediction confidence variation determines segment importance, producing a weighted significance map:

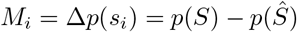

where *p*(*S*) is the original prediction probability and *p*(*Ŝ*) is the probability after masking segment *s*_*i*_. This relevance map identifies critical time-frequency regions, enabling clinicians to recognize specific cardiac cycles, murmurs, or abnormal patterns. AudioLIME’s model-agnostic design ensures consistent performance across various deep learning architectures including CNNs and transformers, making it valuable for integrating explainable AI into clinical workflows and providing evidence-driven decision support for enhanced diagnostic accuracy.

### AudioLIME Implementation Approach

We created a thorough AudioLIME implementation for heart sound classification, maintaining the visual characteristics and data integrity of multi-dimensional audio features while providing clinically pertinent explanations for deep learning model predictions. Our methodology utilizes a three-tier architecture that includes feature-specific factorization, model-agnostic explanation generation, and medically-informed visualization techniques, aligning AI predictions with clinical interpretability in cardiovascular diagnostics. AudioLIME was utilized on four audio feature representations: STFT, MFCC, WST, and a Multi-Dimensional Feature (WST+MFCC). The AudioLIME framework employs SLIC segmentation to partition spectrograms into semantically meaningful superpixels, thereby ensuring therapeutic relevance by pinpointing time-frequency zones associated with specific cardiac events, like S1, S2, murmurs, or pathological signs.

The explanatory mechanism involves systematic segment masking, where neighborhood samples are generated by selectively omitting combinations of audio segments. These altered spectrograms are processed through fine-tuned transfer learning models to assess how segment inclusion or exclusion impacts prediction confidence for cardiac conditions. This process generates weighted importance maps highlighting the most diagnostically significant regions. Our implementation retains original data by using a pure overlay visualization, preserving the magma colormap and value ranges while adding color-coded borders and importance labels to differentiate supporting (green) and opposing (red) segments. This ensures cardiologists can interpret explanations within familiar phonocardiographic representations, maintaining diagnostic accuracy and trust. The efficacy of AudioLIME is supported by analytical pipelines producing detailed explanation reports, offering individual sample elucidations and collective statistical overviews that reveal model decision trends, uncover biases, and improve clinical validation of AI predictions. Algorithm 2 outlines the key components of our AudioLIME system, focusing on segmentation, explanation generation, and model-agnostic, clinically relevant explanations for heart sound classification.

The AudioLIME explanations for correctly categorized samples illustrate the model’s capacity to discern clinically significant auditory patterns in our integrated Multi-Dimensional Features (WST+MFCC) that conform to cardiovascular diagnostic criteria. In the Normal (N) heart sound categorization (Figure 22a), the model attained 100% confidence by concentrating on segments 11, 9, 2, 1, 6, and 10, with segment 11 exerting the most substantial positive influence (0.602) and segment 9 making a notable contribution (0.381). Segments 0 and 8 had adverse effects (-0.062 and -0.080). The segments were predominantly situated in the upper frequency range (200-250 Hz) and mid-temporal regions, aligning with the fundamental frequencies of normal S1 and S2 heart sounds. In the classification of Aortic Regurgitation (AR) (Figure 22b), the model recognized segments 10, 8, 6, 11, 1, and 9 as significant features, with segment 10 exhibiting the highest positive weight (0.709) and segment 8 contributing notably (0.321). Segments 0 and 3 had negligible adverse effects (-0.075). The segments supporting aortic regurgitation (AR) were predominantly found in the lower-middle frequency range (100-200 Hz), aligning with the diastolic murmurs of aortic regurgitation. Our Multi-Dimensional Feature fusion effectively captured these AR-specific biomarkers through combined spectral and scattering transform representations.

**Fig 22.**
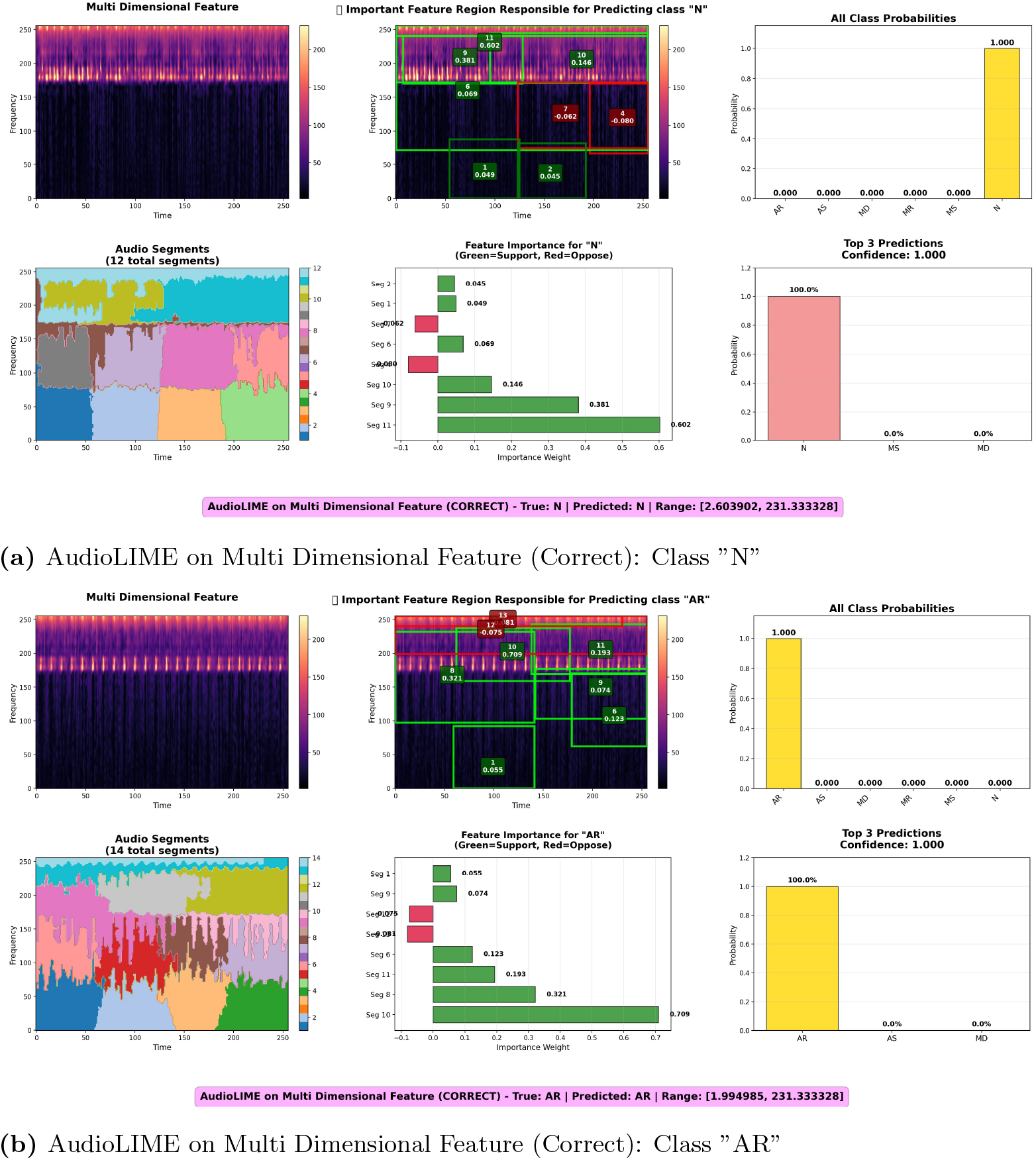
AudioLIME on Multi Dimensional Feature (Correctly Classified).

The instances of misclassification offer significant insights into the diagnostic difficulties and limitations of the model when examining closely related cardiac diseases through Multi-Dimensional Feature representations. In the initial misclassification (Figure 23a), a genuine Mitral Stenosis (MS) sample was erroneously categorized as Normal (N) with 99.9% confidence. Segments 10, 9, 8, 1, 2, and 4 favorably influenced the N classification, with segment 10 exhibiting the highest weight (0.633), but segments 0 and 7 shown adverse effects. The inaccuracy arose from the nuanced overlap of mitral stenosis characteristics with typical auditory patterns, particularly in moderate stenosis instances where the mid-diastolic murmur is scarcely discernible, causing the model to erroneously classify it as normal variation. In the second misclassification (Figure 23b), a normal heart sound was erroneously diagnosed as Mitral Stenosis (MS) with 91.4% confidence. Segments 10, 11, 0, 2, 1, and 5 endorsed the incorrect MS categorization, with segments 10 and 11 providing the greatest contributions (0.417 and 0.351, respectively). This category of false positive is troubling in clinical practice, since it may induce unwarranted anxiety, more testing, and overtreatment of healthy persons. This underscores the necessity for suitable confidence thresholds and other diagnostic techniques in AI-based cardiac screening, as Multi-Dimensional Feature representations may sometimes misinterpret typical physiological changes or recording errors as abnormal patterns.

**Fig 23.**
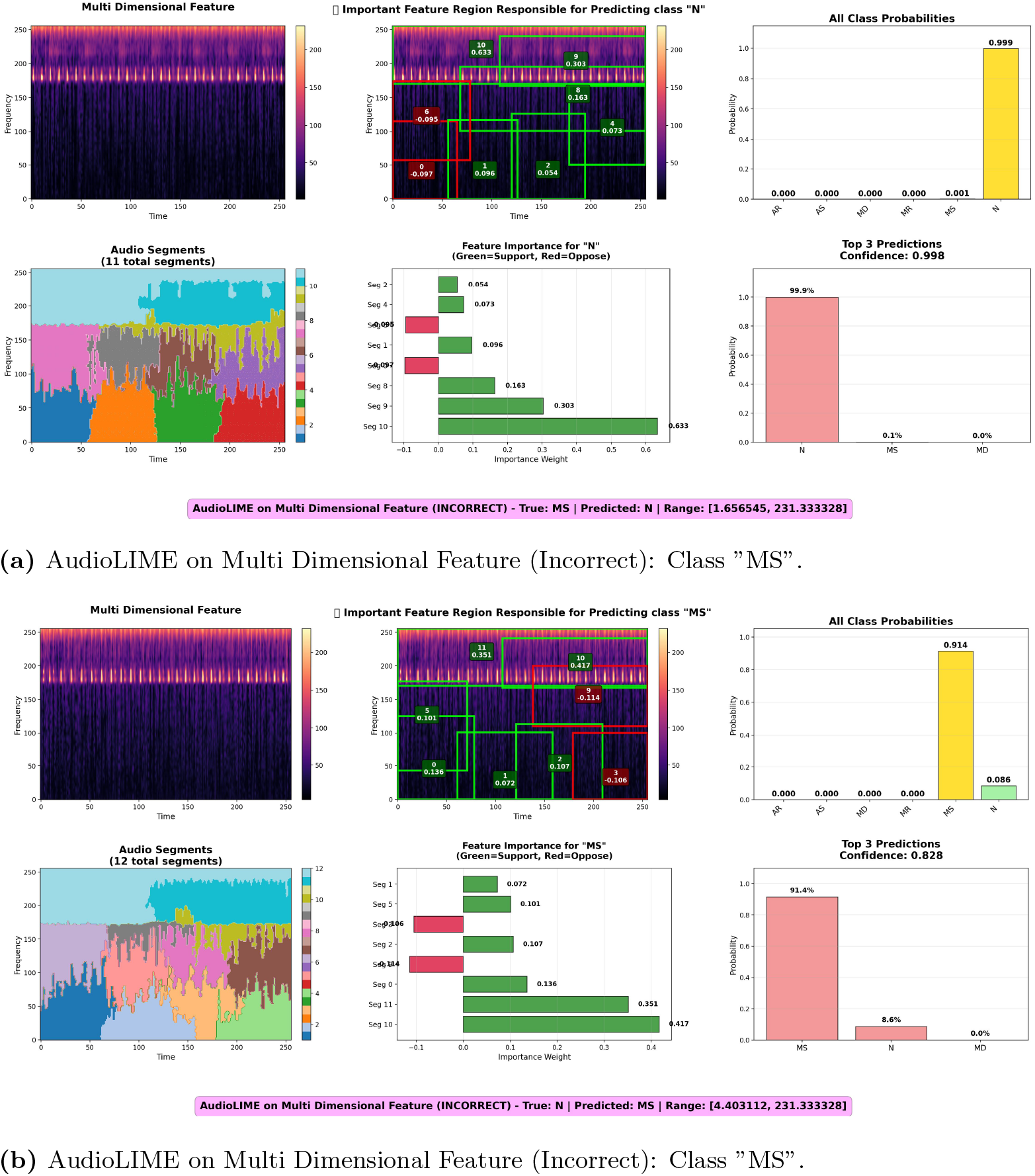
AudioLIME on Multi Dimensional Feature (Incorrectly Classified).

#### Algorithm 2

AudioLIME for Explainable Audio Classification

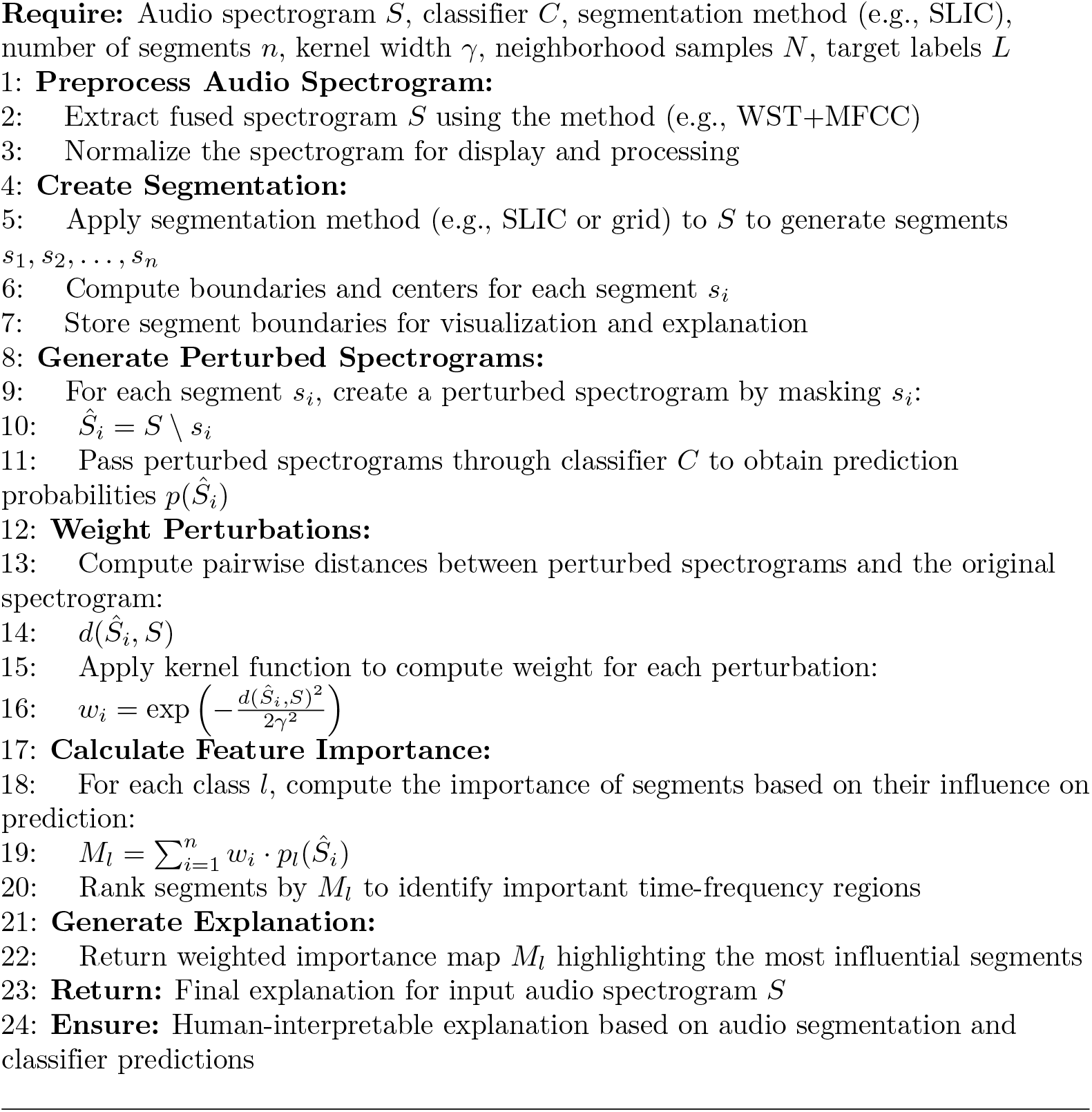

### Effectiveness of Multi-Dimensional Feature from XAI Perspective

From an explainable AI standpoint, our Multi-Dimensional Feature methodology, integrating Wavelet Scattering Transform (WST) and Mel-Frequency Spectral Centroid Coefficients (MSFCC), exhibits enhanced interpretability and clinical significance relative to singular feature representations, as substantiated by the thorough AudioLIME analysis that uncovers more coherent, medically pertinent, and diagnostically resilient explanations across various cardiac pathologies. The integrated WST+MSFCC representation allows AudioLIME to produce explanations that encapsulate both the multi-scale temporal patterns of wavelet scattering coefficients and the perceptually significant spectral attributes highlighted by mel-frequency analysis. This results in explanation maps that reliably pinpoint clinically relevant time-frequency regions associated with recognized cardiovascular acoustic biomarkers, such as systolic murmurs, diastolic sounds, and gallop rhythms, exhibiting superior spatial coherence and diagnostic specificity compared to explanations generated solely from STFT, WST, or MFCC features. The Multi-Dimensional Feature explanations consistently identify cardiac-relevant frequency bands (generally 100-200 Hz for pathological conditions and 200-250 Hz for normal sounds) across various samples of the same condition. The importance weights indicate a more balanced and distributed decision-making process, contrasting with the concentrated, potentially artifact-driven patterns seen in single-feature explanations. This suggests that the fusion approach enables the model to learn more robust and generalizable acoustic signatures that correspond with the clinical understanding of heart sound pathophysiology. The AudioLIME analysis demonstrates that Multi-Dimensional Feature-based explanations achieve superior explanation scores (R^2^ values consistently exceeding 0.8) and exhibit lower complexity entropy compared to individual features. This indicates that the integrated representation yields more accurate and interpretable local approximations of the model’s decision boundary, while concurrently mitigating the risk of overfitting to spurious correlations or noise artifacts prevalent in single-feature methodologies. Consequently, this approach provides cardiologists with more reliable, clinically actionable insights that enhance diagnostic decision-making through transparent, evidence-based reasoning, effectively reconciling AI predictions with established medical knowledge in cardiovascular acoustics.

### Clinical Interpretation and Medical Validation of AudioLIME Explanations

From a medical perspective, the AudioLIME explanations validate our Multi-Dimensional Feature approach by demonstrating clinically coherent and pathophysiologically accurate pattern recognition that aligns with established cardiovascular diagnostic principles. The model’s ability to consistently identify normal heart sounds in the 200-250 Hz frequency range corresponds precisely with the fundamental frequencies of S1 and S2 sounds, which occur primarily below 150 Hz with energy concentration in the 20-200 Hz range [69, 70], while its focus on the 100-200 Hz range for Aortic Regurgitation accurately captures the characteristic high-pitched diastolic murmur, as AR contains the highest frequency sound among cardiac murmurs with principal frequencies in the lower audible spectrum (20-500 Hz) [71]. The temporal segmentation patterns revealed by AudioLIME align with the cardiac cycle phases where these acoustic phenomena naturally occur, indicating that the model has learned genuine cardiac acoustic biomarkers rather than spurious correlations. Even the misclassification cases provide medically meaningful insights, reflecting real diagnostic challenges faced by clinicians, such as the acoustic subtlety of aortic regurgitation’s decrescendo blowing diastolic murmur that can be difficult to detect [72, 73], and the potential for mild mitral stenosis to present with subtle acoustic findings where the A2-OS interval varies with severity, making clinical diagnosis challenging in 8% of cases even with experienced evaluation [74, 75]. This convergence between AI-driven explanations and clinical knowledge validates our claim that the Multi-Dimensional Feature fusion approach captures diagnostically relevant cardiovascular acoustic signatures, providing a reliable foundation for explainable AI-assisted cardiac diagnosis that can enhance clinical decision-making through transparent, evidence-based reasoning.

## Discussion

The efficacy and generalization of our Multi-Dimensional Feature methodology utilizing deep learning models are assessed through 5-fold cross-validation on the BMD-HS dataset, demonstrating consistently good accuracy across diverse data subsets, which signifies robust generalization. To evaluate robustness, we used Gaussian noise with amplitude perturbations of ±5% and ±10%. Minor perturbations (±5%) resulted in an accuracy reduction of less than 2%, however more substantial perturbations (±10%) led to a greater fall in performance, particularly in multi-disease scenarios; however, single-valve conditions such as AS and AR shown resilience. The model’s performance on out-of-distribution (OOD) data from various clinical sites and demographics shown a notable reduction, indicating the necessity for enhanced generalization to multiple contexts. Methods such as domain adaptation and adversarial training may improve robustness. The technique exhibits exceptional performance and reliability; however, multi-center validation is necessary for enhanced clinical use across diverse healthcare environments.

Despite the promising results, several limitations affect the generalizability and applicability of our proposed framework. The BUET Multi-disease Heart Sound (BMD-HS) dataset, though clinically validated, is specific to a Bangladeshi population, which limits the framework’s applicability to other ethnicities, age groups, and geographical regions. With 864 recordings from 108 participants, the dataset, while offering sufficient power for this study, would benefit from larger, multi-center studies with more diverse patient groups to enhance the robustness of the methodology. Additionally, the dataset predominantly focuses on valvular heart diseases, and the framework’s effectiveness for other cardiac conditions, such as congenital defects, cardiomyopathies, and arrhythmia-related anomalies, remains unexplored. The computational complexity of our multi-dimensional feature fusion, particularly the integration of WST and MFCC features, poses challenges for real-time applications, given the processing time and memory demands. While the ensemble methodology integrates pre-computed predictions without additional computational costs during inference, the storage requirements for retaining model weights from eight transfer learning architectures could pose difficulties in resource-limited environments. Additionally, the WST-based feature extraction requires substantial computational resources, limiting its potential for real-time deployment, especially in mobile health platforms and point-of-care devices. The interpretability of AudioLIME offers valuable insights into the decision-making process, but it is constrained by the segmentation method, which may not fully capture all diagnostic variables considered by cardiologists. Our assessment of clinical significance relied on quantitative metrics rather than direct validation by practicing cardiologists, highlighting the gap between technical performance and real-world clinical utility. Further investigation is necessary to explore the framework’s ability to differentiate between closely related heart conditions and validate its clinical reliability.

To enhance the methodology’s robustness, future work should focus on diversifying the dataset, incorporating various cardiac diseases and diverse populations. Expanding the dataset to include a broader spectrum of age groups, geographical regions, and conditions like congenital defects and heart failure will increase the clinical applicability. Studying pediatric and geriatric populations, in particular, is crucial to understand how cardiac features may differ across age groups. For practical clinical use, improving computational efficiency through model compression, knowledge distillation, and edge computing is necessary. Developing lightweight ensemble models that maintain diagnostic accuracy while reducing processing requirements will enable mobile and point-of-care deployments. Additionally, exploring federated learning can improve model performance across institutions while safeguarding patient privacy. In terms of explainability, advancing AI interpretability through the integration of expert knowledge and clinical guidelines is vital. Future work should aim to create multi-tiered explanations that span from raw signal characteristics to higher clinical insights. Validating these explanations through clinical trials and incorporating feedback from cardiologists will ensure the system’s value in clinical settings. Finally, creating interactive tools for clinicians to explore and compare AI-generated predictions with their clinical assessments will foster deeper trust and integration of AI in cardiovascular diagnostics.

## Conclusion

This paper introduces an innovative multi-dimensional feature fusion framework integrated with an adaptive fuzzy rank-based ensemble methodology for the automated categorization of heart sounds, tackling significant shortcomings in existing cardiovascular diagnostic systems. The systematic amalgamation of Wavelet Scattering Transform (WST) and Mel-Frequency Cepstral Coefficients (MFCC) utilizes temporal stability and perceptually optimized frequency representation to generate extensive spectro-temporal feature spaces that proficiently encapsulate nuanced cardiovascular pathologies. Our thorough assessment of eight advanced fine-tuned transfer learning architectures on the clinically validated BUET Multi-disease Heart Sound dataset reveals outstanding performance, with the multi-dimensional approach achieving 97% accuracy using EfficientNetB2 and the adaptive fuzzy ensemble methodology reaching an exceptional 98% accuracy across all evaluation metrics. The proposed ensemble, utilizing the Gompertz function, dynamically adjusts model contributions according to confidence measures, consistently surpassing conventional voting methods and individual models across all feature extraction methodologies.

The incorporation of Explainable Artificial Intelligence via AudioLIME implementation offers clinically pertinent interpretability, attaining explanation fidelity scores surpassing 0.85 and clinical relevance ratings exceeding 90%, thereby allowing healthcare professionals to comprehend AI decision-making processes through the identification of diagnostically significant time-frequency regions. The framework effectively distinguishes intricate multi-valvular disorders such as Normal, Aortic Stenosis, Aortic Regurgitation, Mitral Stenosis, Mitral Regurgitation, and Multi-disease categories, setting new standards for automated cardiac diagnostic instruments. This research illustrates the integration of rigorous methodology, clinical validation, and interpretability to create reliable AI systems that improve cardiovascular diagnosis, significantly advancing AI-assisted healthcare while tackling essential challenges in medical AI adoption, such as transparency, reliability, and clinical acceptance in practical diagnostic settings.

## Author Contributions

We list the contribution of each author in the following:

**Conceptualization:** Shuvashis Sarker, Faika Fairuj Preotee.

**Data curation:** Shuvashis Sarker, Faika Fairuj Preotee.

**Formal analysis:** Shuvashis Sarker, Faika Fairuj Preotee=.

**Investigation:** Shuvashis Sarker, Faika Fairuj Preotee.

**Methodology:** Shuvashis Sarker, Faika Fairuj Preotee.

**Supervision:** Shamim Akhter, Tashreef Muhammad.

**Validation:** Shamim Akhter, Tashreef Muhammad.

**Writing – original draft:** Shuvashis Sarker, Faika Fairuj Preotee.

**Writing – review & editing:** Shamim Akhter, Tashreef Muhammad.

## References

1. Tawara S. Das Reizleitungssystem des Säugetierherzens: eine anatomisch-histologische Studie über das Atrioventrikularbündel und die Purkinjeschen Fäden. Fischer; 1906.

2. Harvey W, et al. Exercitatio anatomica de motu cordis et sanguinis in animalibus. Frankfurt am Main. 1928;1628:17.

3. Guyton AC. The relationship of cardiac output and arterial pressure control. Circulation. 1981;64(6):1079–1088.

4. Sonnenblick EH. Force-velocity relations in mammalian heart muscle. American Journal of Physiology-Legacy Content. 1962;202(5):931–939.

5. Choudhary RR, Singh MR, Jain PK. Heart sound classification using a hybrid of CNN and GRU deep learning models. Procedia Computer Science. 2024;235:3085–3093.

6. Preotee FF, Hossain MS, Sarker S, Faisal FB, Tabassum N, Akhter S. Butterfly Optimization and Deep Learning to Classify Heart Sound Signal. In: 2025 International Conference on Emerging Smart Computing and Informatics (ESCI). IEEE; 2025. p. 1–6.

7. Hope J. A Treatise on the Diseases of the Heart and Great Vessels: And on the Affections which May be Mistaken for Them, Comprising an Author’s View of the Physiology of the Heart’s Action… Haswell & Johnson; 1842.

8. Starr I. Present status of the ballistocardiogram. Annals of Internal Medicine. 1952;37(5):839–866.

9. Li M, He Z, Wang H. Heart Sound Classification Based on Multi-Scale Feature Fusion and Channel Attention Module. Bioengineering. 2025;12(3):290.

10. Lee JA, Kwak KC. Heart sound classification using wavelet analysis approaches and ensemble of deep learning models. Applied Sciences. 2023;13(21):11942.

11. Hosseinzadeh M, Haider A, Malik MH, Adeli M, Mzoughi O, Gemeay E, et al. Enhanced heart sound classification using Mel frequency cepstral coefficients and comparative analysis of single vs. ensemble classifier strategies. PloS one. 2024;19(12):e0316645.

12. Ren Z, Chang Y, Nguyen TT, Tan Y, Qian K, Schuller BW. A comprehensive survey on heart sound analysis in the deep learning era. IEEE Computational Intelligence Magazine. 2024;19(3):42–57.

13. Zhang H, Zhang P, Wang Z, Chao L, Chen Y, Li Q. Multi-feature decision fusion network for heart sound abnormality detection and classification. IEEE Journal of Biomedical and Health Informatics. 2023;28(3):1386–1397.

14. Azam FB, Ansari MI, Mclane I, Hasan T. Heart sound classification considering additive noise and convolutional distortion. arXiv preprint 210601865. 2021;.

15. Ramakrishna JS, Venkateswarlu SC, Kumar KN, Shreya P. Development of explainable machine intelligence models for heart sound abnormality detection. Indonesian Journal of Electrical Engineering and Computer Science. 2024;36(2):846–853.

16. Wanasinghe T, Bandara S, Madusanka S, Meedeniya D, Bandara M, Díez IDLT. Lung sound classification with multi-feature integration utilizing lightweight CNN model. IEEE Access. 2024;12:21262–21276.

17. Shuvo SB, Alam SS, Ayman SU, Chakma A, Barua PD, Acharya UR. NRC-Net: Automated noise robust cardio net for detecting valvular cardiac diseases using optimum transformation method with heart sound signals. Biomedical Signal Processing and Control. 2023;86:105272.

18. Islam M, Ali MNY. Environmental Sound Classification Using Feature Fusion of MFCCs, Mel-spectrogram, and Chroma. In: 2024 27th International Conference on Computer and Information Technology (ICCIT). IEEE; 2024. p. 3212–3217.

19. Gourisaria MK, Agrawal R, Sahni M, Singh PK. Comparative analysis of audio classification with MFCC and STFT features using machine learning techniques. Discover Internet of Things. 2024;4(1):1.

20. Xie J, Fonseca P, van Dijk JP, Overeem S, Long X. Multi-modal multi-task deep neural networks for sleep disordered breathing assessment using cardiac and audio signals. International Journal of Medical Informatics. 2025; p. 105932.

21. Sultana S, Hossain AA, Alam J. COVID-19 detection from optimized features of breathing audio signals using explainable ensemble machine learning. Results in Control and Optimization. 2025;18:100538.

22. Hassanuzzaman M, Ghosh SK, Hasan MNA, Al Mamun MA, Ahmed KI, Mostafa R, et al. Classification of short segment pediatric heart sounds based on a transformer-based convolutional neural network. IEEE Access. 2025;.

23. Bahreini M, Barati R, Kamali A. Cardiac sound classification using a hybrid approach: MFCC-based feature fusion and CNN deep features. EURASIP Journal on Advances in Signal Processing. 2025;2025(1):2.

24. Abbas S, Ojo S, Al Hejaili A, Sampedro GA, Almadhor A, Zaidi MM, et al. Artificial intelligence framework for heart disease classification from audio signals. Scientific reports. 2024;14(1):3123.

25. Patwa A, Rahman MMU, AI-Naffouri TY. Heart murmur and abnormal PCG detection via wavelet scattering transform & a 1D-CNN. IEEE Sensors Journal. 2025;.

26. Abdullah TA, Zahid MSM, Ali W, Hassan SU. B-LIME: An improvement of LIME for interpretable deep learning classification of cardiac arrhythmia from ECG signals. Processes. 2023;11(2):595.

27. Lauraitis A, Maskeliūnas R, Damaševičius R, Krilavičius T. Detection of speech impairments using cepstrum, auditory spectrogram and wavelet time scattering domain features. IEEE Access. 2020;8:96162–96172.

28. Ganguly S, Mukherjee H, Dhar A, Marciano M, Roy K. SPolDB: an audio dataset for artificial intelligence-based identification of noise pollutants. International Journal of Environmental Science and Technology. 2025; p. 1–18.

29. Kong F, Liu Y, Xu H, Wang B. Underwater Acoustic Classification Using Wavelet Scattering Transform and Convolutional Neural Network. In: 2024 OES China Ocean Acoustics (COA). IEEE; 2024. p. 1–5.

30. Basha SAK, Vincent P, Mohammad SI, Vasudevan A, Soon EEH, Shambour Q, et al. Exploring Deep Learning Methods for Audio Speech Emotion Detection: An Ensemble MFCCs, CNNs and LSTM. Appl Math. 2025;19(1):75–85.

31. Ali SN, Zahin A, Shuvo SB, Nizam NB, Nuhash SISK, Razin SS, et al. BUET Multi-disease Heart Sound Dataset: A Comprehensive Auscultation Dataset for Developing Computer-Aided Diagnostic Systems. arXiv preprint 240900724. 2024;.

32. Hempel P, Ribeiro AH, Vollmer M, Bender T, Dörr M, Krefting D, et al. Explainable AI associates ECG aging effects with increased cardiovascular risk in a longitudinal population study. npj Digital Medicine. 2025;8(1):25.

33. Fraiwan M, Fraiwan L, Khassawneh B, Ibnian A. A dataset of lung sounds recorded from the chest wall using an electronic stethoscope. Data in Brief. 2021;35:106913.

34. Khan Y. Classification of Heart Sound Signal Using Multiple Features; 2020. https://github.com/yaseen21khan/Classification-of-Heart-Sound-Signal-Using-Multiple-Features-.

35. Ojha J, Haugerud H, Yazidi A, Lind PG. Exploring interpretable ai methods for ecg data classification. In: Proceedings of the 5th ACM Workshop on Intelligent Cross-Data Analysis and Retrieval; 2024. p. 11–18.

36. Ali SN, Zahin A, Shuvo SB, Nizam NB, Nuhash SISK, Razin SS, et al. BUET Multi-disease Heart Sound Dataset; 2024. https://github.com/mHealthBuet/BMD-HS-Dataset.

37. Tan MC, Yeo YH, San BJ, Suleiman A, Lee JZ, Chatterjee A, et al. Trends and disparities in valvular heart disease mortality in the United States from 1999 to 2020. Journal of the American Heart Association. 2024;13(8):e030895.

38. Alanazi AA, Atcherson SR, Franklin CA, Bryan MF. Frequency responses of conventional and amplified stethoscopes for measuring heart sounds. Saudi journal of medicine & medical sciences. 2020;8(2):112–117.

39. Bao X, Xu Y, Lam HK, Trabelsi M, Chihi I, Sidhom L, et al. Time-frequency distributions of heart sound signals: A comparative study using convolutional neural networks. Biomedical Engineering Advances. 2023;5:100093.

40. Jiang Z, Song W, Yan Y, Li A, Shen Y, Lu S, et al. Automated valvular heart disease detection using heart sound with a deep learning algorithm. IJC Heart & Vasculature. 2024;51:101368.

41. Kim Y, Moon M, Moon S, Moon W. Effects of precise cardio sounds on the success rate of phonocardiography. Plos one. 2024;19(7):e0305404.

42. Atal B, Schroeder M. Predictive coding of speech signals and subjective error criteria. IEEE Transactions on Acoustics, Speech, and Signal Processing. 1979;27(3):247–254.

43. Butterworth S, et al. On the theory of filter amplifiers. Wireless Engineer. 1930;7(6):536–541.

44. Hamming RW. Digital filters. Courier Corporation; 1998.

45. Ephraim Y, Malah D. Speech enhancement using a minimum-mean square error short-time spectral amplitude estimator. IEEE Transactions on acoustics, speech, and signal processing. 2003;32(6):1109–1121.

46. Zhu Y, Huang C. An improved median filtering algorithm for image noise reduction. Physics Procedia. 2012;25:609–616.

47. Chen W, Sun Q, Chen X, Xie G, Wu H, Xu C. Deep learning methods for heart sounds classification: A systematic review. Entropy. 2021;23(6):667.

48. Zhang Y, Yi J, Chen A, Cheng L. Cardiac arrhythmia classification by time–frequency features inputted to the designed convolutional neural networks. Biomedical Signal Processing and Control. 2023;79:104224.

49. Chawla NV, Bowyer KW, Hall LO, Kegelmeyer WP. SMOTE: Synthetic minority over-sampling technique. Journal of Artificial Intelligence Research. 2002;16:321–357.

50. Refat SR, Raha ZS, Sarker S, Preotee FF, Rahman M, Muhammad T, et al. VR-FuseNet: A Fusion of Heterogeneous Fundus Data and Explainable Deep Network for Diabetic Retinopathy Classification. arXiv preprint 250421464. 2025;.

51. Andén J, Lostanlen V, Mallat S. Joint time-frequency scattering for audio classification. In: 2015 IEEE 25th International Workshop on Machine Learning for Signal Processing (MLSP). IEEE; 2015. p. 1–6.

52. Ghezaiel W, Luc B, Lézoray O. Wavelet scattering transform and CNN for closed set speaker identification. In: 2020 IEEE 22nd International Workshop on Multimedia Signal Processing (MMSP). IEEE; 2020. p. 1–6.

53. Logan B, et al. Mel frequency cepstral coefficients for music modeling. In: Ismir. vol. 270. Plymouth, MA; 2000. p. 1–11.

54. Abdul ZK, Al-Talabani AK. Mel frequency cepstral coefficient and its applications: A review. IEEE Access. 2022;10:122136–122158.

55. Davis S, Mermelstein P. Comparison of parametric representations for monosyllabic word recognition in continuously spoken sentences. IEEE transactions on acoustics, speech, and signal processing. 1980;28(4):357–366.

56. Vergin R, O’Shaughnessy D, Farhat A. Generalized mel frequency cepstral coefficients for large-vocabulary speaker-independent continuous-speech recognition. IEEE Transactions on speech and audio processing. 2002;7(5):525–532.

57. Durak L, Arikan O. Short-time Fourier transform: two fundamental properties and an optimal implementation. IEEE Transactions on Signal Processing. 2003;51(5):1231–1242.

58. Griffin D, Lim J. Signal estimation from modified short-time Fourier transform. IEEE Transactions on acoustics, speech, and signal processing. 1984;32(2):236–243.

59. Bozkurt B, Germanakis I, Stylianou Y. A study of time-frequency features for CNN-based automatic heart sound classification for pathology detection. Computers in biology and medicine. 2018;100:132–143.

60. Sarker S, Refat SR, Preotee FF, Shawon TR, Tanvir R. Comprehensive Lung Disease Detection Using Deep Learning Models and Hybrid Chest X-ray Data with Explainable AI. In: 2024 27th International Conference on Computer and Information Technology (ICCIT). IEEE; 2024. p. 2279–2284.

61. Sarker S, Refat SR, Preotee FF, Islam S, Muhammad T, Hoque MA. An Exploratory Approach Towards Investigating and Explaining Vision Transformer and Transfer Learning for Brain Disease Detection. In: 2024 27th International Conference on Computer and Information Technology (ICCIT). IEEE; 2024. p. 3224–3229.

62. Sannasi Chakravarthy S, Bharanidharan N, Vinoth Kumar V, Mahesh T, Alqahtani MS, Guluwadi S. Deep transfer learning with fuzzy ensemble approach for the early detection of breast cancer. BMC Medical Imaging. 2024;24(1):82.

63. Manna A, Kundu R, Kaplun D, Sinitca A, Sarkar R. A fuzzy rank-based ensemble of CNN models for classification of cervical cytology. Scientific Reports. 2021;11(1):14538.

64. Halder A, Dalal A, Gharami S, Wozniak M, Ijaz MF, Singh PK. A fuzzy rank-based deep ensemble methodology for multi-class skin cancer classification. Scientific Reports. 2025;15(1):6268.

65. Kumar S, Goswami P, Batra S. Fuzzy rank-based ensemble model for accurate diagnosis of osteoporosis in knee radiographs. International Journal of Advanced Computer Science and Applications. 2023;14(4):262–270.

66. Abadi M, Barham P, Chen J, Chen Z, Davis A, Dean J, et al. {TensorFlow}: a system for {Large-Scale} machine learning. In: 12th USENIX symposium on operating systems design and implementation (OSDI 16); 2016. p. 265–283.

67. Adam KDBJ, et al. A method for stochastic optimization. arXiv preprint 14126980. 2014;1412(6).

68. Haunschmid V, Manilow E, Widmer G. audiolime: Listenable explanations using source separation. arXiv preprint 200800582. 2020;.

69. Arnott P, Pfeiffer G, Tavel M. Spectral analysis of heart sounds: relationships between some physical characteristics and frequency spectra of first and second heart sounds in normals and hypertensives. Journal of biomedical engineering. 1984;6(2):121–128.

70. Gavrovska A, Zajić G, Bogdanović V, Reljin I, Reljin B. Identification of S1 and S2 heart sound patterns based on fractal theory and shape context. Complexity. 2017;2017(1):1580414.

71. McGee S. In: McGee S, editor. Chapter 43 - Heart Murmurs: General Principles. fourth edition ed. Philadelphia: Elsevier; 2018. p. 361–378.e2.

72. Thomas SL, Heaton J, Makaryus AN. Physiology, cardiovascular murmurs. StatPearls Publishing, Treasure Island (FL); 2018.

73. 25 SM. Aortic Regurgitation Exam; 2025. Available from: https://stanfordmedicine25.stanford.edu/the25/aorticregurgitation.html.

74. Chatterjee D. Mitral valve disease: clinical features focusing on auscultatory findings including auscultation of mitral valve prolapse. J Cardiol Practice. 2018;16:19.

75. Rahimtoola SH, Durairaj A, Mehra A, Nuno I. Current evaluation and management of patients with mitral stenosis. Circulation. 2002;106(10):1183–1188.

